# Evolutionary history and rhizosphere microbial community composition in domesticated hops (*Humulus lupulus L*.)

**DOI:** 10.1101/2024.10.30.621085

**Authors:** Alexandra McElwee-Adame, Raya Esplin-Stout, Trevor Mugoya, George Vourlitis, Nautica Welch, Kayser Afram, Maryam Ahmadi Jeshvaghane, Nathan Bingham, Alexis Dockter, Jacob Eslava, Giovanni Gil, Joshua Mergens, Amran Mohamed, Tram Nguyen, Fatum Noor, Nathan Salcedo, Arun Sethuraman

## Abstract

*Humulus lupulus L.*, commonly known as hops, is a perennial crop grown worldwide and is well known for its pharmacological, commercial, and most importantly brewing applications. For hundreds of years, hops have undergone intense artificial selection with over 250 cultivated varieties being developed worldwide, all displaying differences in key characteristics such as bitter acid concentrations, flavor and aroma profiles, changes in photoperiod, growth, and pathogen/pest resistances. Previous studies have individually explored differences between cultivars, aiming to identify markers that can quickly and cost-effectively differentiate between cultivars. However, little is known about their evolutionary history and the variability in their associated rhizospheric microbial communities. Coupling phenotypic, genomic, and soil metagenomic data, our study aims to explore the global population structure and domestication history of 98 hops cultivars. Additionally, we assessed differences in growth rates, rates of viral infection, usage of dissolvable nitrogen, and soil microbial community compositions between US and non-US based cultivars. Contrary to previous studies, our study revealed that worldwide hop cultivars cluster into four primary subpopulations; Central European, English, and American ancestry as previously reported, and one new group, the Nobles, revealing further substructure amongst Central European cultivars. Modeling the evolutionary history of domesticated hops reveals an early divergence of the common ancestors of modern US cultivars around 2800 ybp, and more recent divergences with gene flow across English, Central European, and Noble cultivars, reconciled with key events in human history and migrations. Furthermore, cultivars of US origin were shown to overall outperform non-US cultivars in both growth rates and usage of dissolvable nitrogen and display novel microbial composition.

## Introduction

Domestication and repeated selective breeding of plants by humans has opened new horizons beyond human history, unleashing novel bursts of molecular evolutionary dynamics and selection for agronomic traits within domesticated plant genomes, spanning across trophic levels. Recent studies of domesticated plant genomes have revealed signatures of selection for pungency, shape, and size in peppers (Liu et al., 2023), heterosis through introgression and the spread of advantageous alleles for shattering, grain size, and disease/pathogen resistance in rice (reviewed in Chen et al., 2019) and for kernel size and number in corn (Yang et al., 2023), auto- and allo-polyploidization, and pangenomic evolution of chromatin structures and non-coding regulatory variation in cotton (Wang et al., 2022), post-transcriptional transposon-mediated regulation of yield-related traits in maize (Sun et al., 2023), and reduced genomic diversity due to serial bottlenecks in a variety of crop species (reviewed in Alam and Purugganan 2024). Meanwhile, the examination of trophic-level interactions and coevolutionary dynamics during domestication include examples in human and other domesticator evolutionary trajectories (summarized in Jackson 1996, Purugganan 2022). Similarly, rhizospheric microbial and fungal communities have been shown to coevolve in response to domestication in rice (Chang et al., 2021), apples (Abdelfattah et al., 2021), lima beans (da Silva et al., 2022), bread wheat (Yue et al., 2023), soybeans (Luo et al., 2022) and tomatoes (Smulders et al., 2021). Understanding the complex interplay between genomics of domestication with phenotypic and ecosystem dynamics is key to developing agronomically robust crops that are resilient to novel thermal regimes, soil types, and emerging pathogens.

*Humulus lupulus L.*, commonly known as hops, is a dioecious perennial crop that natively grows in Asia, Europe and North America (Jakse et al., 2004). Found in the family Cannabaceae, *H. lupulus* diverged from its closest relative, *Cannabis sativa*, roughly 27.8 million years ago (McPartland, 2018). The provenance of origin of *H. lupulus* is unknown, however it is believed to have originated in China, similar to *Cannabis* and its sister species *H. japonicus* and *H. yunnanensis* that are only distributed in the Asiatic region (Ren et al., 2021; Small, 1978). Five subspecies of *H. lupulus* have been described, with two subspecies naturally occurring in Europe and Asia (var. *lupulus* and var. *cordifolius*) and the remaining three subspecies distributed across North America (var. *pubescens* in the midwest and eastern United States, var. *lupuloides* in Canada and eastern US, and var. *neomexicanus* in the Western US (Reeves & Richards, 2011; Small, 1978).

Hops are a prominent cash crop that is utilized extensively in the brewing industry, relying on bitter alpha (humulones) and beta (lupulones) acids that are produced by hop lupulin glands within female cones. These unique bittering compounds impart the necessary bitterness in beer, often used to offset the sweetness produced by the grain of choice. Today, there are over 250 hop cultivars developed and domesticated worldwide, each differing in bitter acid concentrations, essential oil composition, vigor/growth, and disease tolerance (Castro 2021), with the United States and Germany being the top producers worldwide (Acosta-Rangel et al. 2024). Previous phylogenetic analyses conducted on wild hops using ETS sequences and cpDNA markers suggest that wild European hops diverged from Asian and North American hop subspecies around 1.05-1.27 MYA with Asian and North American hop subspecies diverging again about 0.46-0.69 MYA (Murakami et al., 2006). Additionally, previous population genetics analyses utilizing microsatellite markers suggest a significant differentiation between European and North American hops with significant internal differentiation between cultivated and wild populations within the same regions (Stajner et al., 2008). With advancements in biotechnology and reducing costs of genomic sequencing, new studies have begun to focus efforts into genome assembly and gene discovery in hops, in an effort to improve crop production (Bolger et al., 2014; Edwards & Batley, 2010; Kumar et al., 2021).

For instance, recent efforts to genetically fingerprint hop cultivars with SNP and SSR markers from repositories and germplasms across the United States by Driskill et al., 2022 revealed (a) high clonality and relatedness among cultivars that are often cultivated/marketed under different names, (b) geographical population structure that separates cultivars into Wild North American (and USA-developed), English, and Continental European subpopulations, and (c) significant degrees of admixture (as estimated using SSR markers) across all domesticated and wild cultivars. With the new availability of a high-quality phased genome of the Cascade hop (Padgitt-Cobb et al., 2023), and recent efforts to develop genome-wide SNP markers (John Henning, pers. comm.), we sought to answer several outstanding questions in hop biology: (1) what is the ancient and contemporary evolutionary history of domestication in hops, (2) can we reconcile genomic data with historical records of cultivation and spread in the species?, (3) can we quantify dynamics and differences amongst rhizospheric microbial communities across domesticated hop cultivars of different provenances?, and (4) do cultivars of different provenance exhibit phenotypic variation in ecologically and agronomically relevant traits such as growth rate, leaf surface area, soil moisture content, hop-virus and mildew resistance, and pathogen-related leaf damage? We address these questions using a combination of molecular methods to quantify genomic variation across hop cultivars, computational modeling to assess their evolutionary history, and molecular and computational methods from a common-garden greenhouse experiment to assess variability in the ecology and microbial community dynamics of US-based and non-US-based hop cultivars.

## Methods

### Cultivar collection, genomic DNA extractions and QC

Genomic DNA was extracted from 163 hop accessions from the USDA National Clonal Germplasm Repository (NCGR) and subject to genotyping by sequencing (GBS) according to the protocol described in Driskill et al., 2022. Briefly, 30-50 mg of fresh leaf tissue was homogenized using a mill and high molecular weight gDNA was extracted using a modified E-Z 96 Plant DNA extraction kit protocol (Omega BioTek, Norcross, GA, USA). DNA quality and quantity were ascertained using a Tecan Infinite M Plex multimode plate reader (Tecan Group Ltd, Zürich, Switzerland) and diluted to 3 ng/μL, followed by GBS using the ApeK1 enzyme, barcoded, and sequenced on an Illumina HiSeq 3000 according to the protocol described in Elshire et al., 2011, followed by variant calling as described by Driskill et al., 2022. The final SNP dataset comprising 143,309 SNPs was then further filtered to remove triploids, biallelic sites (--min-alleles 2 –max-alleles 2) in VCFTools (Danecek et al., 2011). Subsequently we filtered for sites with a minor allele frequency below a threshold of 0.05 to remove monomorphic and other non-informative sites with low degree of polymorphism, potentially due to sequencing and genotyping errors. To ensure thorough removal of all triploids from the vcf file, we compiled a list of known triploid cultivars from hopslist resource (https://www.hopslist.com/) and used vcftools --remove parameter to parse and delete any records if present. The final filtered dataset comprised 27,163 bi-allelic SNPs across 98 cultivars.

### Population Genomic Analyses

Using the filtered vcf file, we explored the population structuring and differentiation amongst our 98 cultivars. To explore the genetic diversity amongst hops cultivars, we used Arlequin (v.3.5.2.2) to determine the number of SNP sites per cultivar, observed/ expected homozygosity, inbreeding coefficient (F-statistic) and observed/expected heterozygosity.

First we converted our filtered vcf file to an Arelquin file using the software PGDSpider v. 2.0 (Lischer and Excoffier 2012) and the converted file analyzed with Arelqeuin (v 3.5.2.2) (Excoffier and Lischer 2010). Degree of relatedness amongst each cultivar was computed using vcftools v.4.2 (Danecek et al., 2011).

To determine population structuring amongst our 98 cultivars, we performed unsupervised clustering analysis using the software ADMIXTURE, with an admixture model with acceleration, testing K=1 to K=10 (Alexander et al. 2009).

To determine the optimal number of subpopulations (here denoted as K), a five-fold cross-validation approach was used. Once the optimal number of cultivar populations was identified, we further explored the population differentiation between subpopulations, estimated as Weir and Cockerham’s F_st_ using Arlequin v.3.5.22 (Excoffier and Lischer 2010). To explore the within population level phylogenetic relationships amongst cultivars, we utilized a maximum likelihood approach employed by IQ-TREE v.2.3.6 (Nguyen et al., 2014). First, the filtered VCF file was converted into NEXUS format using the software PGDSpider v. 2.0 (Lischer and Excoffier 2012), creating a concatenated sequence consisting of SNPs derived from the VCF file. Next, we performed a multiple sequence alignment on concatenated sequences using the software MAFFT v.7.525 (Katoh et al. 2002).

Once our sequences were aligned, IQ-TREE was employed using (-m MFP) to determine the optimal nucleotide substitution model to use during phylogenetic construction and (-bb 10000) to perform an ultrafast bootstrap approximation on the consensus tree with 10,000 replicates. The IQ-TREE run was also parallelized across 20 CPUs (-T 20). The consensus tree was then visualized and annotated using FigTree v.1.4.5 (Rambaut 2009).

### Demographic modeling of evolutionary history

Using the number of subpopulations identified ADMIXTURE (K = 4), we constructed the 2-dimensional derived allele frequency spectra distributions using PPP (Webb et al., 2021). We then employed a combination of a tool we developed - CoalMiner v1.0 (Esplin-Stout and Sethuraman, *in prep*, https://github.com/raywray/CoalMiner) and fastsimcoal28 (Excoffier et al., 2023) to determine the most likely evolutionary model. Using CoalMiner, we randomly generated 1000 different putative evolutionary demographic models based on known parameters about hops, incorporating variation in migration rates, population size variation, admixture rates, and divergence times. Each individual demographic model was run 1000 times in *fastsimcoal*28, with each run consisting of 1000 coalescent simulations. The parameters used in the CoalMiner + fastsimcoal28 pipeline were: uniform per-site mutation rate distribution of min: 6.1e-9, max: 6.1e-9; uniform effective population size distribution of min: 10, max: 2e6; log uniform migration rate distribution of min: 1e-5, max: 5; and a uniform divergence time distribution of min: 1, max: 5000 generations. After collating multiple independent runs of fastsimcoal28, we determined the best-fitting model and corresponding parameters by selecting the model with the smallest difference between maximum observed likelihood of the model (under the observed DAFS distributions) and the maximum estimated likelihood under the simulated model. Thereon, AIC of the best model was calculated using an R script by Vitor Sousa (https://speciationgenomics.github.io/fastsimcoal2/<u>, pers. comm.</u>). Visualization of the best model was generated using an R script by Joana Meier (https://speciationgenomics.github.io/fastsimcoal2/). Confidence intervals around the parameter estimates of the best run were then computed using a 100 replicate parametric bootstrap analysis.

### Greenhouse Experiments

Ten replicate saplings of five US-based cultivars of hops (Brewer’s Gold, Columbus, Neomexicanus, Comet, Zeus) and five non-US-based cultivars (Hallertauer, Southern Cross, Saaz 72, Fuggle, Sorachi Ace) were obtained from Great Lakes Hops (Dutch Touch Growers, Zeeland, MI, USA), with their provenance ascertained by SSR genotyping of gDNA extracted from young leaf material, amplified at 8 microsatellite markers (K910, K221, PI2018, P62011, HIAGA7, K931, K852, K016) using the protocols of Driskill et al., 2022 & Nahla Bassil (pers. comm.). Saplings were transplanted into half-gallon pots and replenished with Miracle Grow potting soil in late June 2023 and maintained on a 2-3 day per week watering schedule depending on weather. Trellises were constructed to permit vines’ ascension about 1 month after transplanting. Plants received no external fertilizer throughout the greenhouse experiment.

### Plant Growth Measurement

Aboveground plant biomass was measured on September 12, 2023 before the fall senescence period. Above ground biomass was harvested, dried at 72°C for approximately 1 week, and weighed on a digital balance. Because the plants were similar in size when transplanted on 12 July, 2023, the change in biomass was used to characterize growth of each cultivar over the ca. 62 day growing period.

### Quantifying soil extractable nitrogen

Dissolved N (NH_4_+NO_3_) was measured colorimetrically on a randomly chosen subset of samples from each cultivar (n = 2 pots/cultivar) from 2M KCl extractions following Mulvaney (1996) for NH_4_ and Miranda et al. (2001) for NO_3_. Briefly, 10g of sieved topsoil from each cultivar was mixed with 40 ml of 2M KCl and shaken for 1 hour on a shaker table. The results were then filtered, and the supernatants were collected and stored in 30 ml plastic bottles. The NH_4_ and NO_3_ concentrations were analyzed using a spectrophotometer at 667 nm for NH_4_ (Mulvaney 1996) and 540 nm for NO_3_ (DeForest 2007; Miranda et al. 2001).

### Quantifying soil moisture

Percent soil moisture was measured gravimetrically as [(M_f_ - M_d_)/M_d_] x 100, where M_f_ = the fresh mass of soil collected from each pot and M_d_ is the dry mass of soil after drying in an oven at 105°C for 1 week.

### Soil DNA extractions, NGS library prep, 16s rRNA and ITS sequencing

Rhizospheric top soil (~2 inches in depth) was obtained from well-established greenhouse plants, along with a “control” soil sample in August 2023 by students of the Fall 2023 BIOL 596 “Research Methods in Agricultural Sciences” at San Diego State University, sealed in sterile bags, and stored immediately at −80C until further processing. Soil microbial and fungal gDNA was extracted using the ZymoBIOMICS 96 MagBead DNA Kit (Zymo Research, Irvine, CA), and quality of the DNA was assessed using agarose gel electrophoresis. DNA quantification was performed on a Qubit Fluorometer 4.0 (Thermo Fisher Scientific) using a broad range fluorescence kit, and shipped to Zymo Research, Irvine, CA for targeted library preparation and sequencing. Briefly, the Quick-16s Primer Set V3-V4 (for microbial community sequencing) was used in rtPCRs, followed by quantification using qPCR. Pooled libraries were then cleaned and concentrated, prior to quantification on a TapeStation (Agilent Technologies, Santa Clara, CA) and a Qubit Fluorometer. Sequencing was then performed on an Illumina NextSeq 2000 with p1 reagent kits (600 cycles) and a 30% PhiX spike-in.

### Bioinformatic analyses of soil genomic data

The Dada2 pipeline (Callahan et al., 2016) was used to process raw reads, identify unique amplicons, and to remove chimeric sequences. Amplicons were then processed using the *uclust* function in QIIME v.1.9.1 (Caporaso et al., 2010), to assign OTU’s against the Zymo Research Database and compute diversity indices (alpha, beta), and to compute significant abundance differences between groups (US-based versus non-US-based cultivar versus control soils). Thereon, we conducted literature surveys to ascertain nitrogen fixing bacterial taxa amongst identified abundant OTUs.

### Quantifying viral damage to leaves

Viral damage to leaves was measured from digital images on September 5, 2023 (about 1 week prior to the last growth measurement) using ImageJ optical software (V1.54i). Approximately 4-6 fully expanded, middle-aged leaves were randomly sampled from each plant and photographed on a white background using a Cannon digital camera. The percentage of each leaf damaged, defined as a part of the leaf that was chlorotic or with abnormal leaf pigmentation, was quantified as the area of the leaf that was visibly damaged (A_d_) divided by the total leaf area (A_T_) x 100. Using imageJ, leaf images were transformed to a false color image to either highly the total leaf area, or by adjusting the threshold, the area of the leaf that was damaged. These false color images were converted to binary (black and white) images for the calculation of total and damaged leaf area. Percent leaf damage was calculated as (A_d_/A_T_) x 100.

### Relative growth rate modeling as a function of abundance of nitrogen fixers

Relative growth rate of all US and Non-US cultivars were modeled as a function of mean dissolved nitrogen content, bacterial genus relative abundance and bacterial nitrogen fixing capability. Average dissolved nitrogen and relative growth rate measures were normalized and scaled per cultivar. Subsequently, relative abundance data from soil 16s rRNA sequencing was used to create a unified data frame with mapped abundance values to each US and Non-US-based cultivar. The relative abundance of each bacterial genus present per hop cultivar was calculated and recorded in the unified data frame. A “yes”or “no” designation was given to each bacterial genus on their known ability to fix nitrogen. A multiple linear regression model with no interaction between predictors was fit using the lm() function in R (R Core Team 2024). To ensure we met the homoscedasticity, correlation between residuals, multicollinearity, independence and normality of residuals assumptions of the multiple linear regression model we evaluated the model response variable with the Breush-Pagan, Durbin Watson, variance inflation factor and autoplot functions respectively (Zeileis & Hothorn 2002, Tang et al. 2016). Model significance was then assessed with *summary* and *anova* functions in R (R Core Team 2024).

The fitted multiple linear regression failed to meet homoscedasticity and normality for the response variable. We further fit a generalized linear model (GLM) with a Poisson distribution with a log link function. Prior to GLM fitting, the average relative growth rate response variable was transformed by adding a positive constant to all values to ensure the data was appropriate for use with a poisson family GLM. The fitted GLM was assessed for overdispersion and determined a GLM with a quasi-poisson distribution was an appropriate fit for the data. We then assessed the resultant model with *summary* and *anova* functions as before.

## Results

### Genetic diversity and differentiation

Average observed genome-wide heterozygosity (Table 1) across all cultivars was estimated to be 0.24, with “Banner” hops having the highest observed heterozygosity (0.43), and “Hersbrucker Alpha” having the lowest heterozygosity (0.09). Genetic differentiation (Weir and Cockerham’s F_st_; Table 2) was lowest between populations sharing Noble Ancestry and Central European ancestry groups and highest between the American and both Noble and Central European ancestry groups.

**Table 1:**
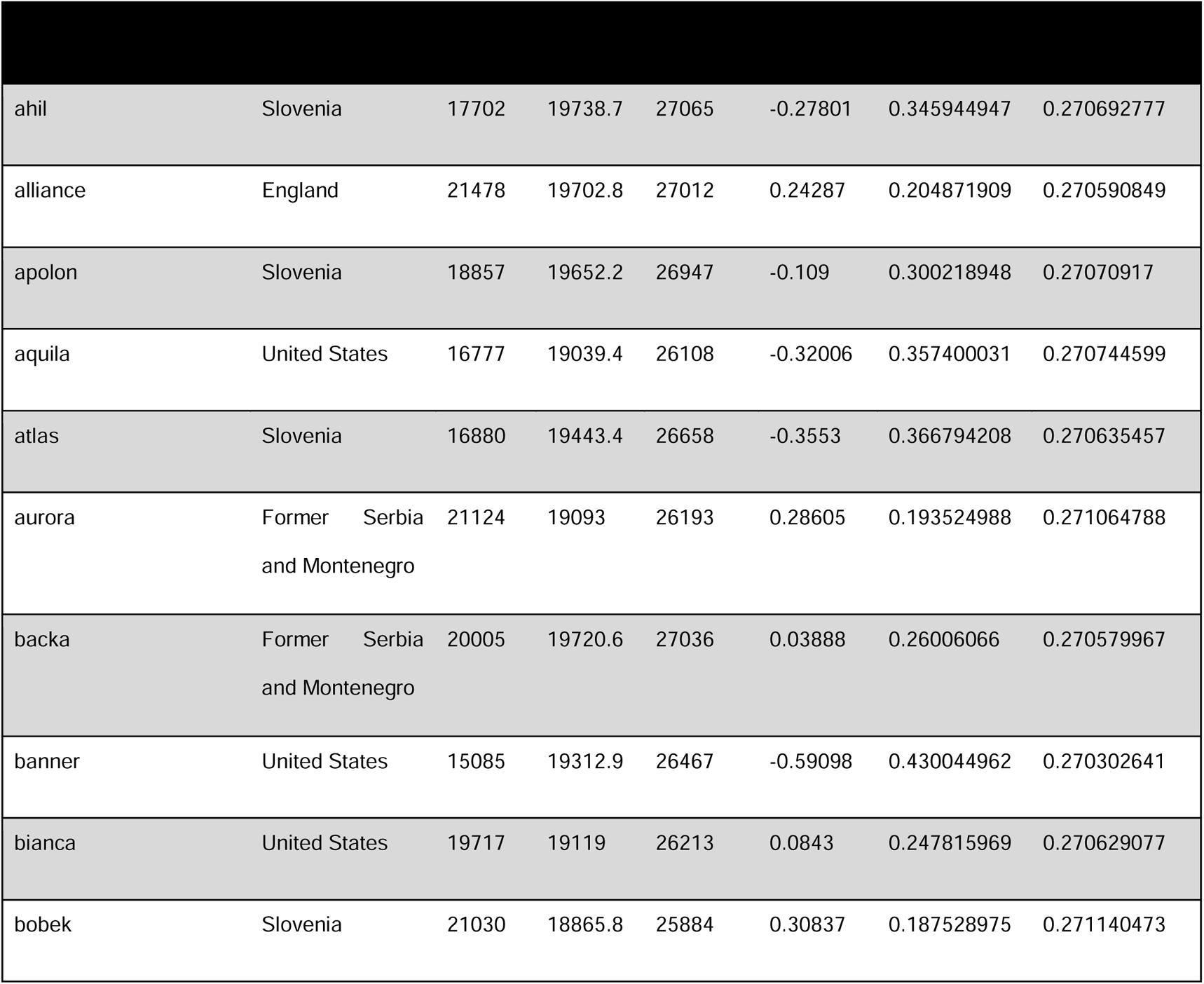
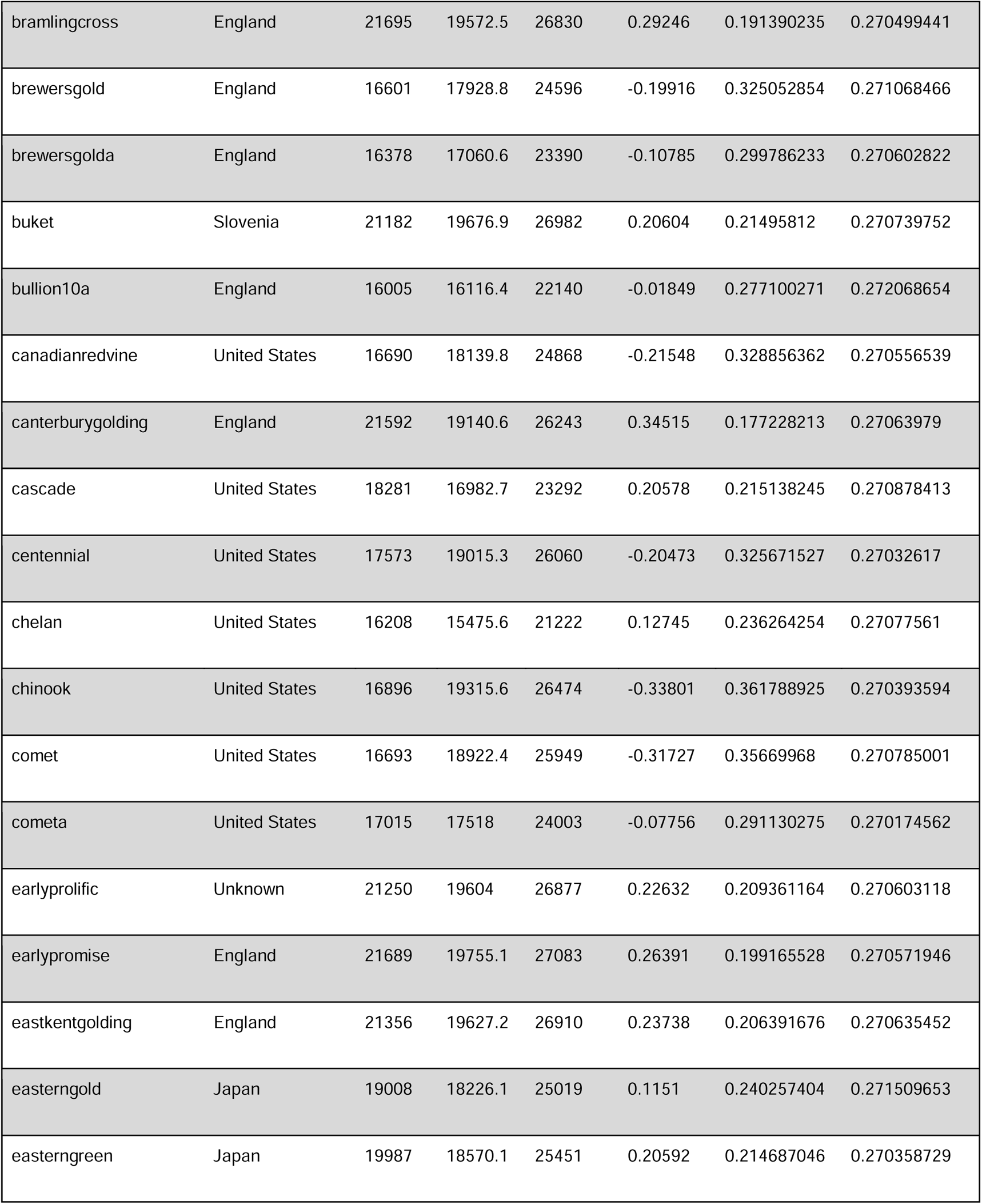
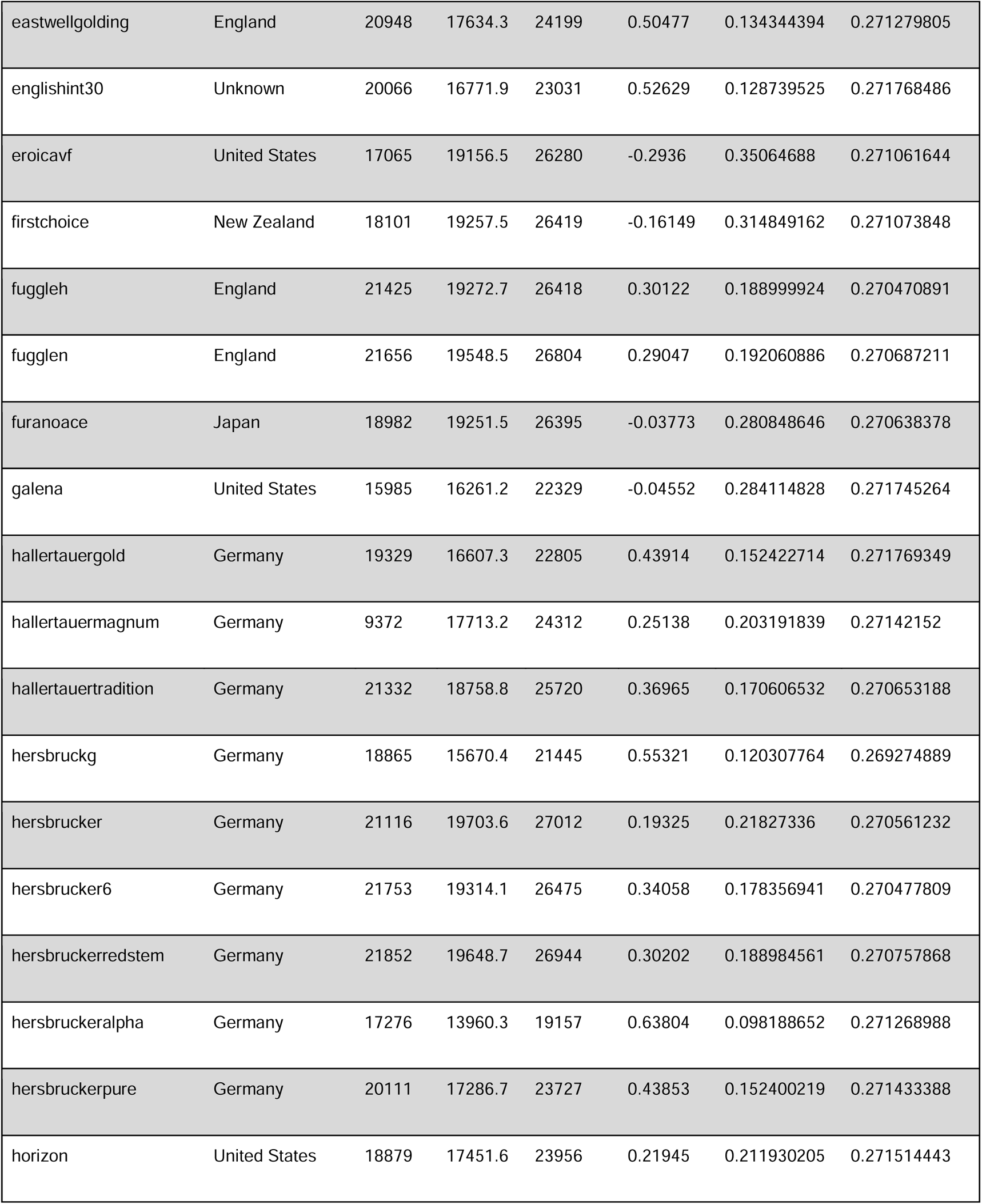
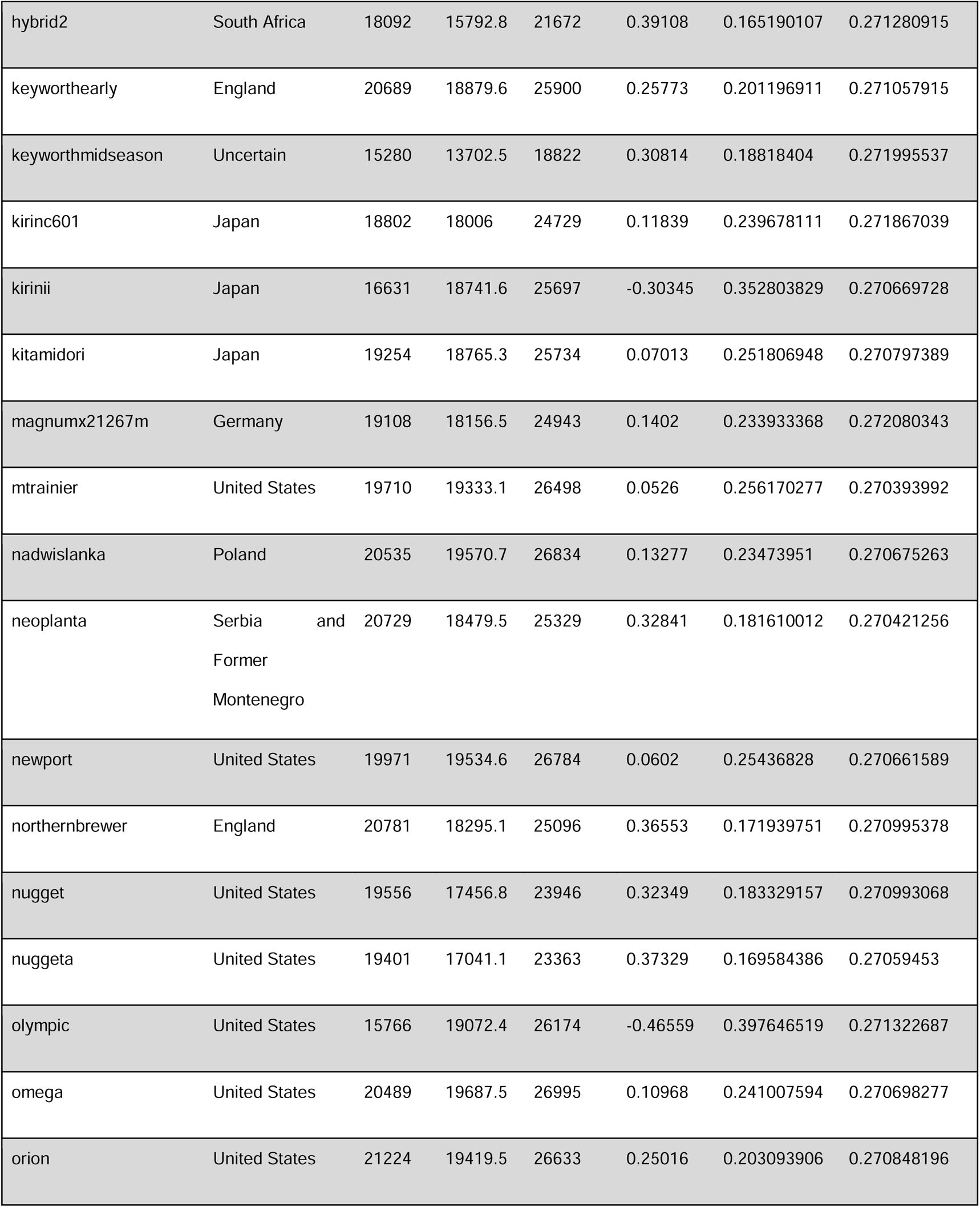
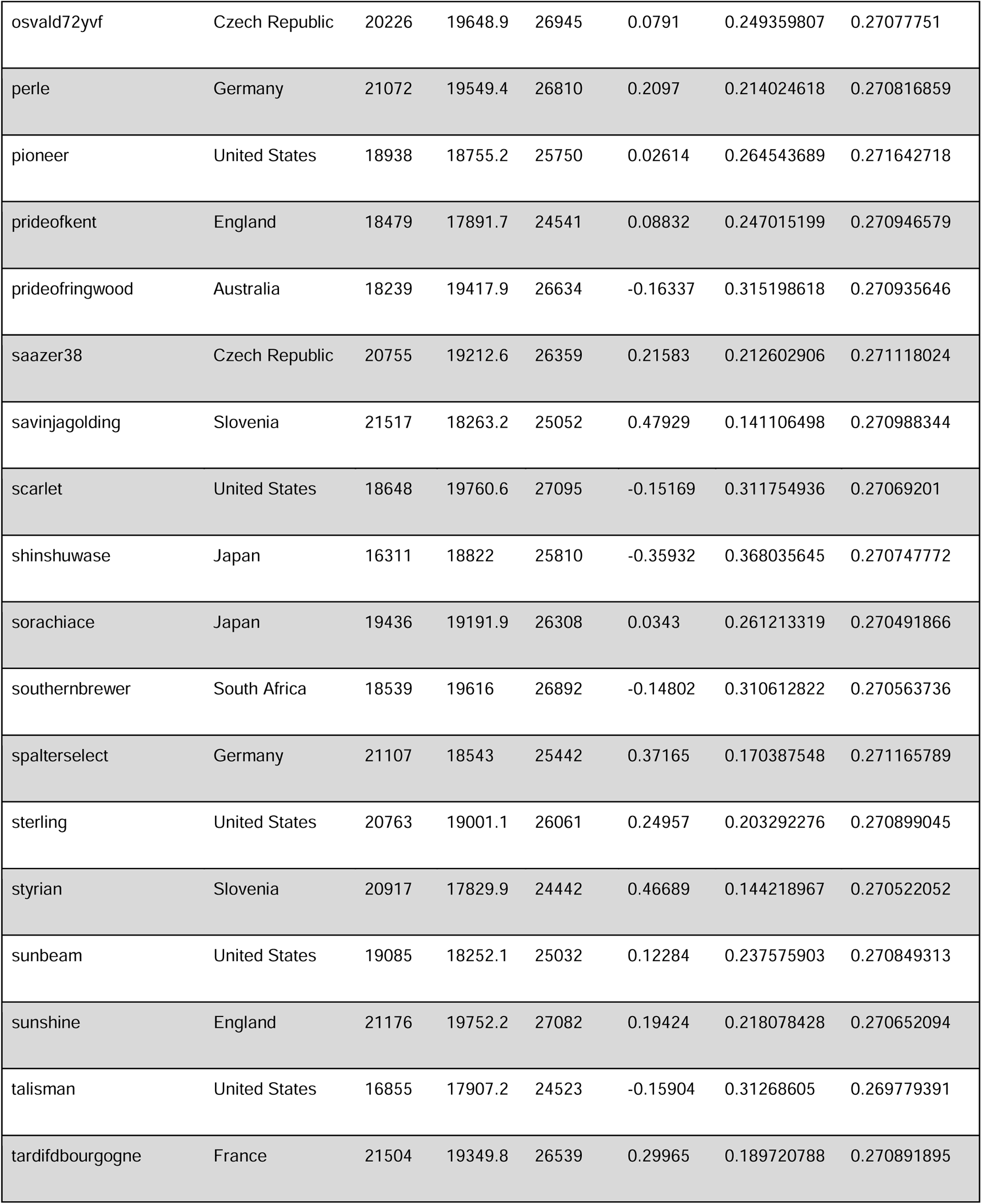
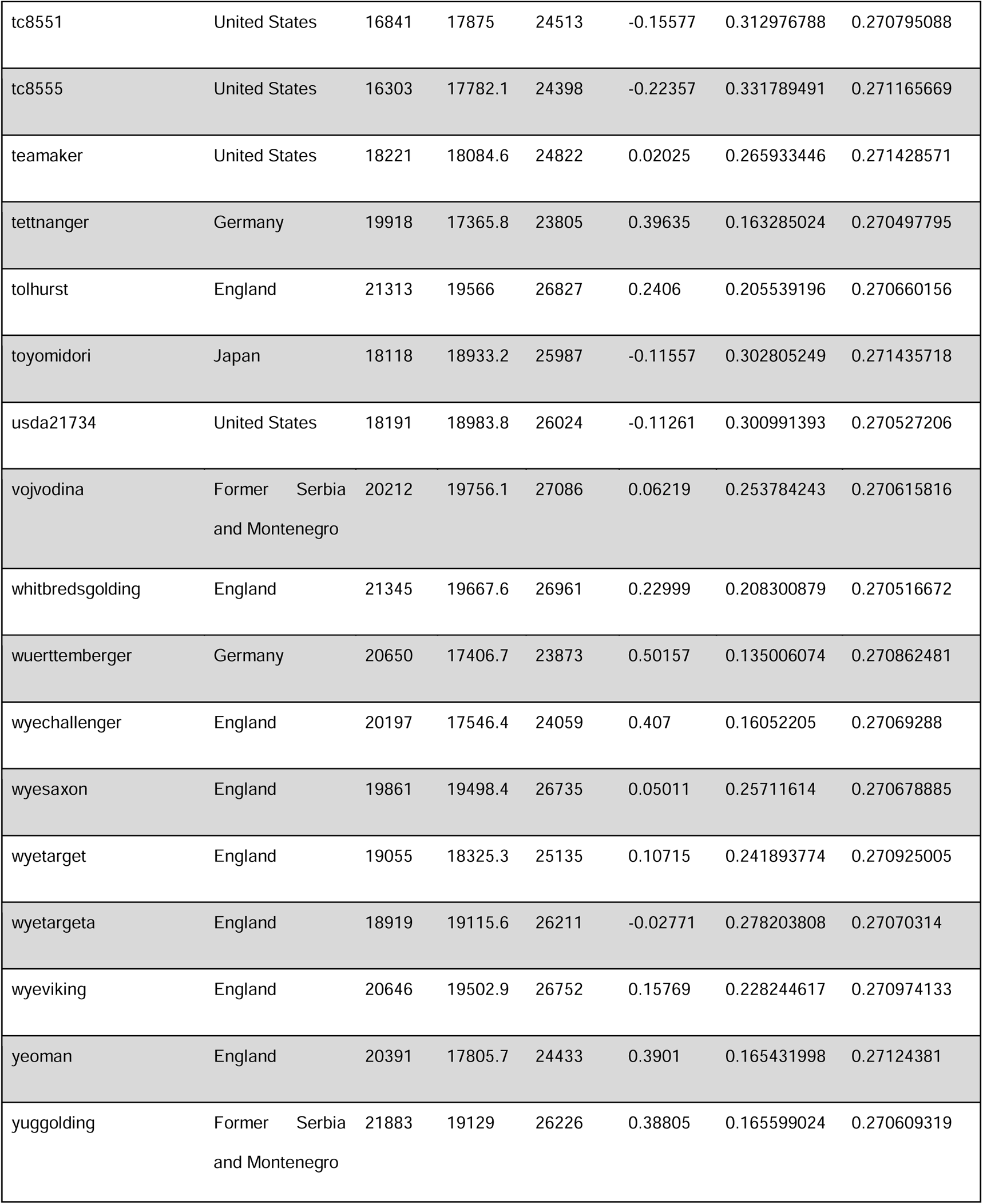
Cultivars included in this study, putative country of origin, observed and expected number of homozygous sites, number of polymorphic sites, F statistic (inbreeding coefficient), and expected and observed heterozygosity as estimated using genome-wide SNPs.

**Table 2:** Estimates of pairwise differentiation between 4 subpopulations of hop cultivars, computed as Weir and Cockerham’s estimator using Arlequin v. 3.5.2.2 (below diagonal) and their corresponding p-values (above diagonal), indicating significant genomic differentiation between all 4 subpopulations.

| Populations | Central European | Noble | English | American |
| --- | --- | --- | --- | --- |
| Central European | - | *** | *** | *** |
| Noble | 0.057654 | - | *** | *** |
| English | 0.11322 | 0.13652 | - | *** |
| American | 0.20043 | 0.2118 | 0.06557 | - |

\*\*\* indicates highly significant values (<0.0005)

### Population Structure and Relatedness

Estimation of population structure using ADMIXTURE shows a high degree of admixture amongst each of the populations, supporting the literature of extensive crossbreeding between cultivars. Using a cross-validation approach, we found the optimal number of subpopulations to be K = 4, with cultivars of Central European ancestry comprising subpopulation Group 1, Noble ancestry for subpopulation Group 2, English ancestry for subpopulation Group 3 and American ancestry for subpopulation Group 4.

We opt here to use the word ‘ancestry’ rather than simply ‘origin’ since while the majority of cultivars found amongst the four population groups also happen to share origins for the same location, this was not the case for all. For instance, the popular cultivar Brewer’s Gold is said to be of English origin due to its development at Wye College in 1919. However, the parental origin of this cultivar is the result of a cross between a wild American hop from Manitoba, Canada and open pollination in England. Genetically, the Brewer’s Gold is largely of American ancestry, but of British origin. This designation was further supported by both our phylogenetic and population structuring results which revealed that Brewer’s Gold is not only dominantly of American ancestry, but also phylogenetically sister to many of its own American progeny (Centennial, Furano Ace, Atlas, Aquila, Olympic, Eroica, Galena and Chelan). Several close relatives were also estimated among analyzed cultivars - Chelan, Galena, Talisman, Banner, Brewer’s Gold, Olympic and Eroica indicated > 0.90 relatedness (hereon, Group A), Kirin II, Shinshuwase, Talisman, Centennial, Olympic, Bullion 10A, Aquila and Chelan with > 0.75 relatedness (hereon, Group B), and Nadwislanka, Saazer 38, Tettnager, Osvald 72 > 0.65 relatedness (hereon, Group C).

### Phylogenetic analyses

Phylogenetic reconstruction using IQ-TREE reflects similar patterns with four major clades corresponding to the four population ancestries as described by the ADMIXTURE analyses. Additionally, our phylogenetic analysis revealed interesting patterns of evolutionary history amongst the four groups. At the largest scale, the first major split to occur amongst the four lineages is between those of Central European ancestry and English/North American ancestry. First, while our ADMIXTURE analysis supports the separation of ‘Noble’ hops cultivars and their progeny into a distinct population group, our phylogenetic analysis revealed that Noble cultivars, while distinct, form a group sister to Central European cultivars. Additionally, while monophyletic clades can be found within the American/English ancestry portion of the tree for their respective populations, overall, both American and English hops form paraphyletic clades, suggesting a complex evolutionary history amongst those of American and English ancestry.

### Complex evolutionary history of domestication

The best supported model reflecting the evolutionary history of cultivated hops using fastsimcoal28 (Excoffier et al., 2020) suggests that the earliest split among domesticated hop cultivars analyzed occurred around 2799 ybp (95% C.I. 2724.41 - 2799.15 ybp) between the common ancestors of modern-day Central European and American cultivars. Post-divergence, there were significant reductions in effective population sizes via bottlenecks, resulting in a contemporary N_e_ of 1415 (95% C.I. 1466.78 - 1511.60) of American cultivars, compared to the common ancestor of all strains with estimated N_e_ of 1,766,785 (95% C.I. of 1,739,913 - 1,767,188). A similar bottleneck and divergence event, occurring nearly 2337 ybp (95% C.I. 2352.95 - 2429.61 ybp) yielded the common ancestors of contemporary Noble cultivars, with an estimated N_e_ of 1376 (95% C.I. 1321.38 - 1365.50). The most recent divergence was estimated to have occurred from the common ancestors of Central European cultivars to establish English strains, around 623.50 ybp (95% C.I. 629.64 - 648.18 ybp). Both Central European and English cultivars are estimated to have comparably much higher N_e_ (> 1,000,000), despite small bottlenecks estimated during their recent history. Significant migration (crossbreeding) events were estimated between contemporary English and Central European cultivars after their recent split. Significant migration was also estimated between Central European and Noble cultivars, with low rates of recent crossing between American and other cultivars.

### Diversity and abundance of microbial communities in US-based versus non-US-based cultivars

Domestic US-based cultivars had an overall lower alpha diversity of bacteria (Fig 5) as well as a lower abundance of nitrogen fixing bacterial taxa in the topsoil compared to foreign (non-US-based) cultivars. Foreign cultivars, on the other hand, had a greater abundance of nitrogen fixing bacteria accompanied by an overall higher alpha diversity of bacteria, which is more than likely attributed to the low selectivity of microorganisms present in foreign cultivar soils. Dissolvable nitrogen levels, which refer to the bioavailable extractable nitrogen in the soils of foreign and domestic cultivars, was higher, but not statistically significant in domestic cultivars when compared to foreign cultivars (p-value = 0.20). Overall relative growth rates using volumetric analysis were significantly higher (p-value < 0.001, F = 4.501, df = 9) in domestic cultivars compared to foreign cultivars. Relative growth rate was positively correlated with dissolvable nitrogen but not genus relative abundance or whether or not the genus is nitrogen fixing and was significantly predicted by the dissolvable nitrogen levels (p-value < 2e-16). Nitrogen is known to play a critical role in vegetative plant growth and hops can remove up to 150 lbs/acre per year, requiring heavy fertilization.

### Multiple Linear Regression Model Assessment

The residuals vs fitted autoplot in addition to the Breusch-Pagan test results show that our model does not conform to the assumption of homoscedasticity (BP = 117.27, df = 5, p-value < 2.2e-16). Independence of model residuals was evaluated with the Durbin-Watson test and showed significant levels of positive autocorrelation in model residuals (DW=0.01379927, p-value = 0). Based on the adjusted generalized variance inflation factor scores for each predictor we found no evidence for multicollinearity between model predictors (Table 2.).

### Poisson Generalized Linear Model Assessment

The residual deviance of the poisson GLM was found to be 59.929 on 221 degrees of freedom which indicates that our model is underdispersed. Additionally, we obtained sufficient evidence to reject the null hypothesis of normality of the distribution of response variable (W = 0.84612, p-value = 3.422e-14).

### Quasi-Poisson Model Fitting

To address the under-dispersion and non-normality of response variable violations of the poisson GLM, a quasi-Poisson was fitted with the log link function. Mean dissolved nitrogen was found to be the only predictor that had any significant influence on mean relative growth rate (t-value = 12.342, p-value < 2e-16). The coefficient would seem to indicate there is a significant positive correlation between average dissolved nitrogen and average relative growth rate (Table 3).

### Biomass production and percent virus damage

There were statistically significant differences in growth between the various cultivars, with Neomexicanus having the highest biomass production and Hallertau MF the lowest (Fig. 3a). There were also significant differences in percent leaf viral damage, however, differences between cultivars were smaller (Fig. 3b). Viral damage was negatively correlated with biomass production (y = 46.1 - 1.59x’ R^2^ = 0.05; p < 0.05; n = 100).

**Fig. 1.**
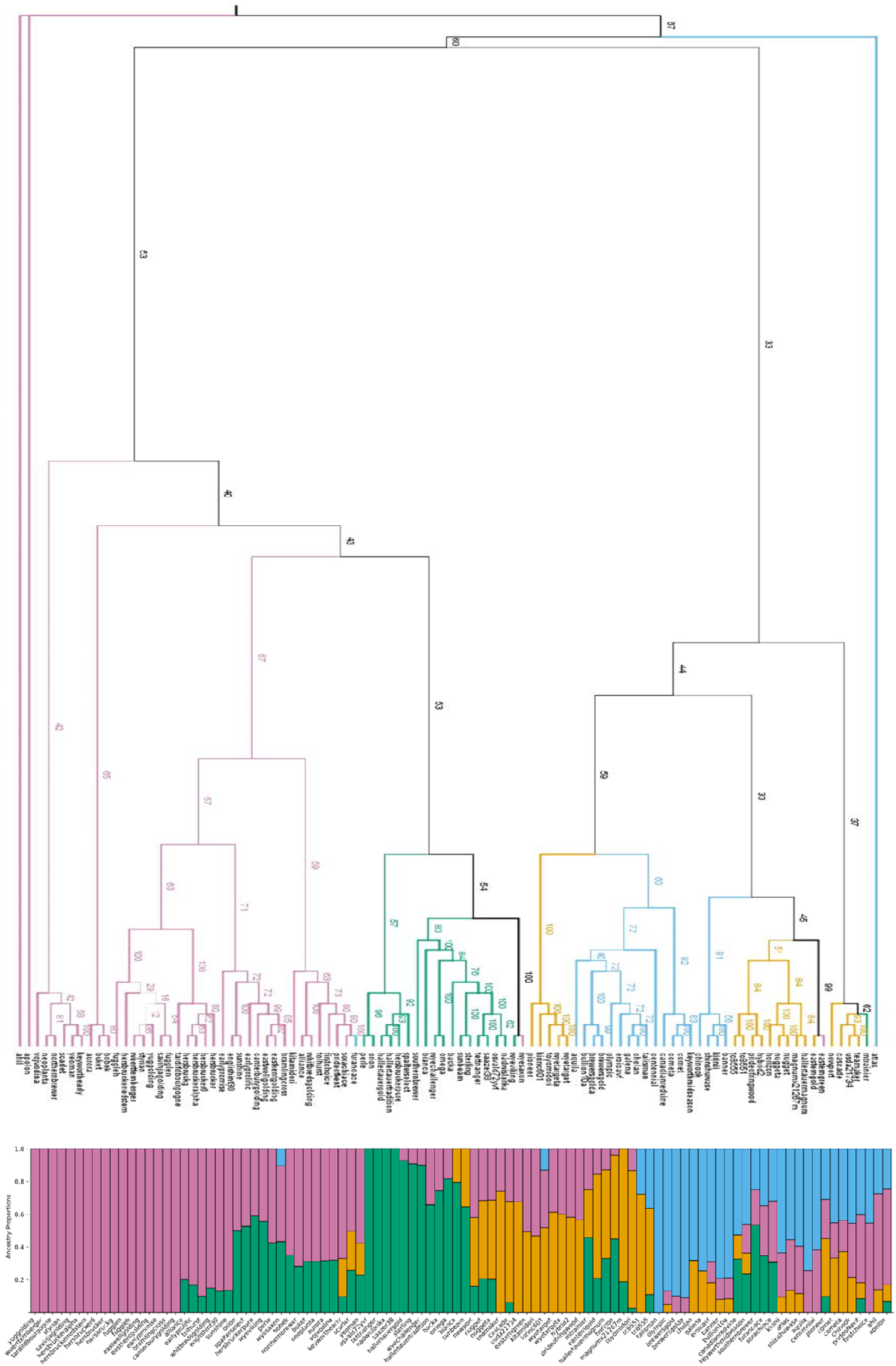
Consensus tree generated from IQ-Tree with bootstrap values along branches (top) and corresponding ancestry proportions of all hop cultivars analyzed using ADMIXTURE (Alexander et al., 2009 - bottom). Cultivars on the tree and stacked bar plots are color coded to indicate their known geographical origins - green indicating “Noble” ancestry, pink indicating hops of “Central European” ancestry, orange denoting “English” ancestry, and light blue representing predominantly “American” ancestry. Note the extensive presence of admixture in domesticated American and English cultivars.

**Fig. 2.**
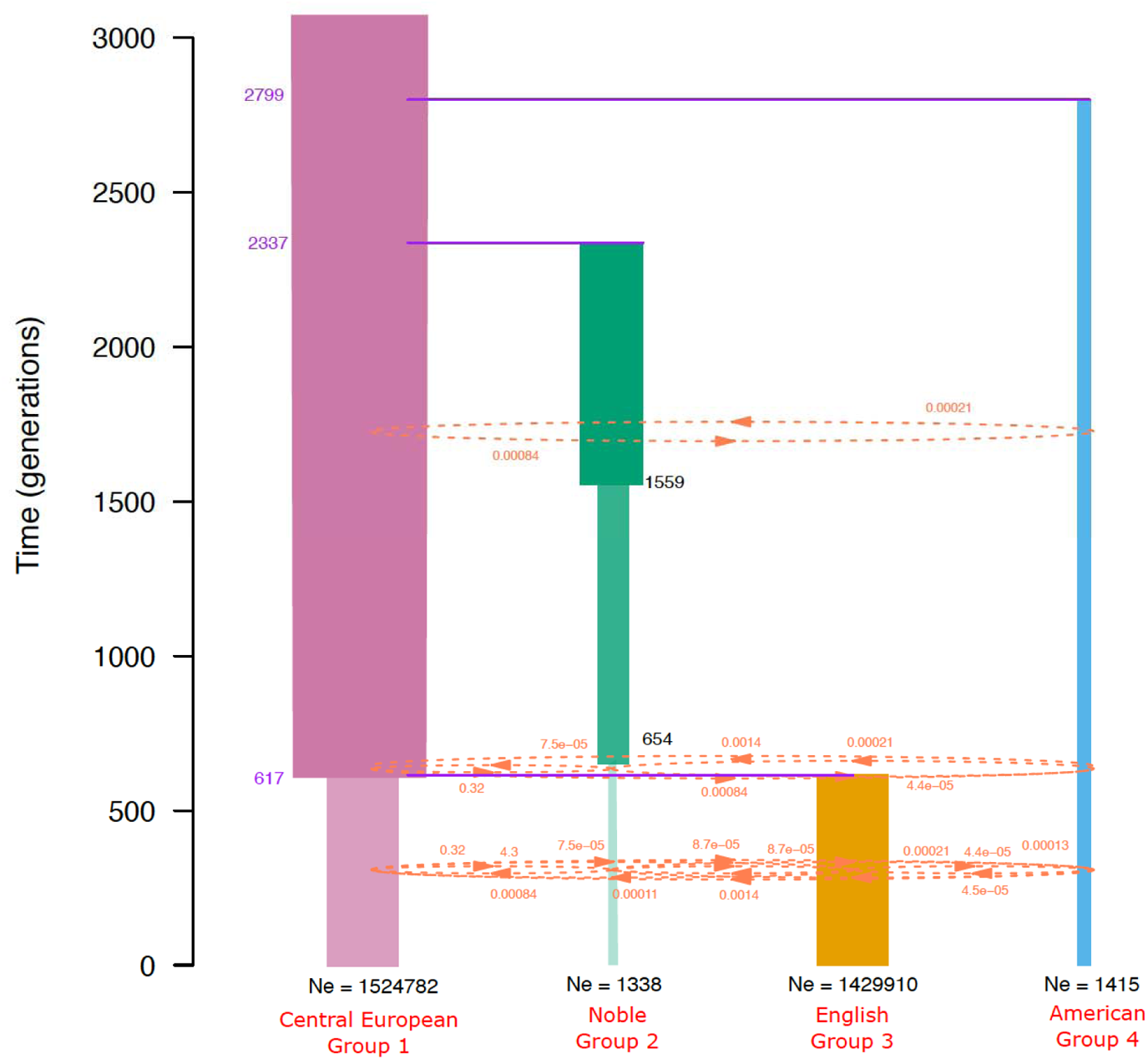
Most likely model of evolutionary history of the four predominant provenances determined by CoalMiner and fastsimcoal28 analyses. Group 1 (pink) = Central European Ancestry, Group 2 (green) = Noble Ancestry, Group 3 (orange) = English Ancestry, Group 4 (light blue) = American Ancestry. Estimates of divergence times are shown in purple font, while estimates of bottleneck times are shown in black. The most likely model also included several significant migration events, here shown in orange dotted lines/arrows.

**Figure 3.**
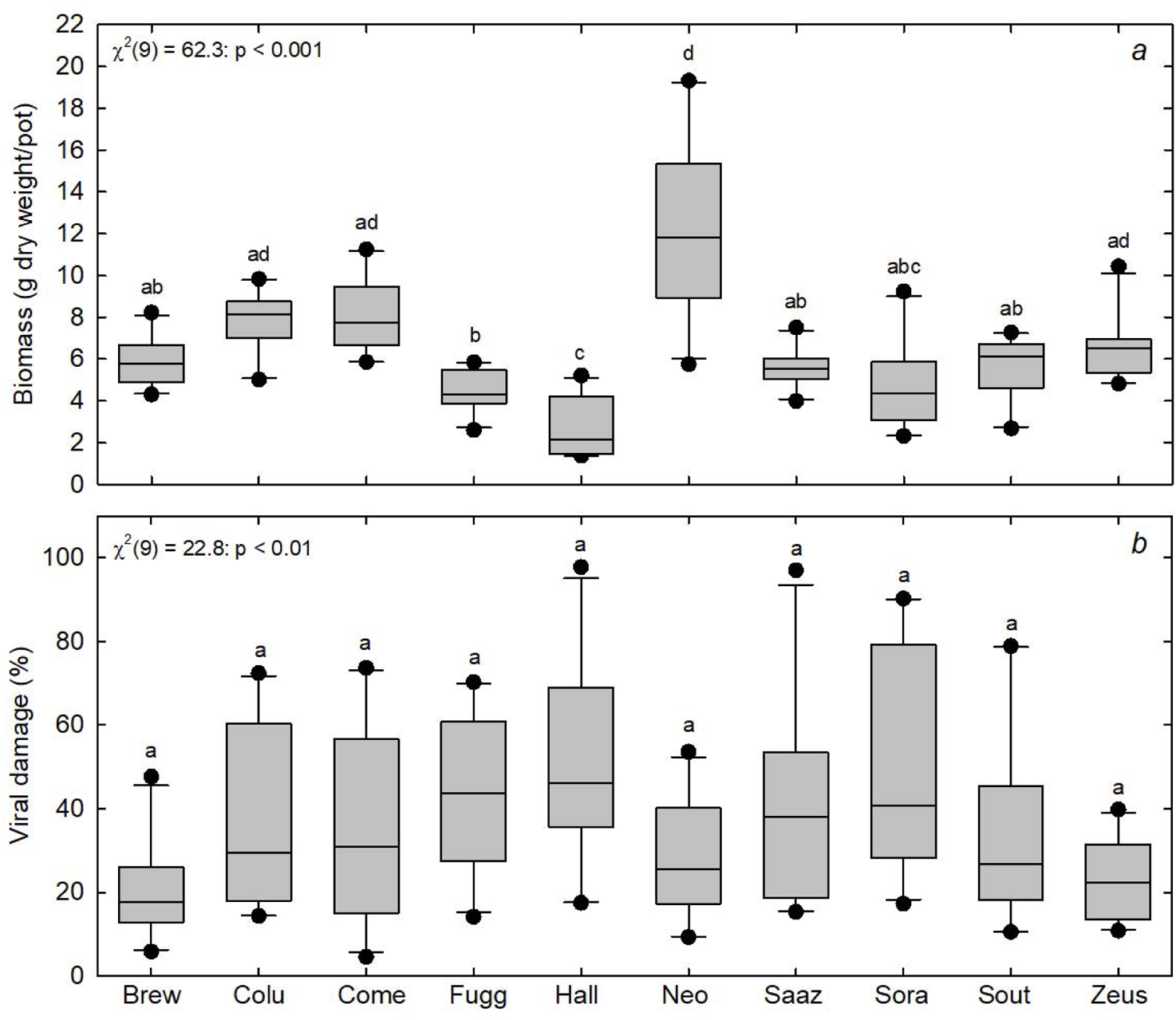
Boxplots of biomass production (a) and leaf viral damage (b) for each cultivar. Statistical differences were assessed using a Kruskal-Wallis ANOVA (*X*^2^ statistic with 9 degrees of freedom) because data were non-normal. Box plots with different lower-cased letters are significantly different according to a Dwass-Steel-Critchlow-Fligner post-hoc pairwise comparison test.

When data were aggregated based on the origin of the cultivar (i.e. US versus non-US). Cultivars that were developed in the US had significantly higher biomass production (Fig. 4a) and lower viral damage (Fig. 4b) than cultivars created outside the US.

**Fig. 4.**
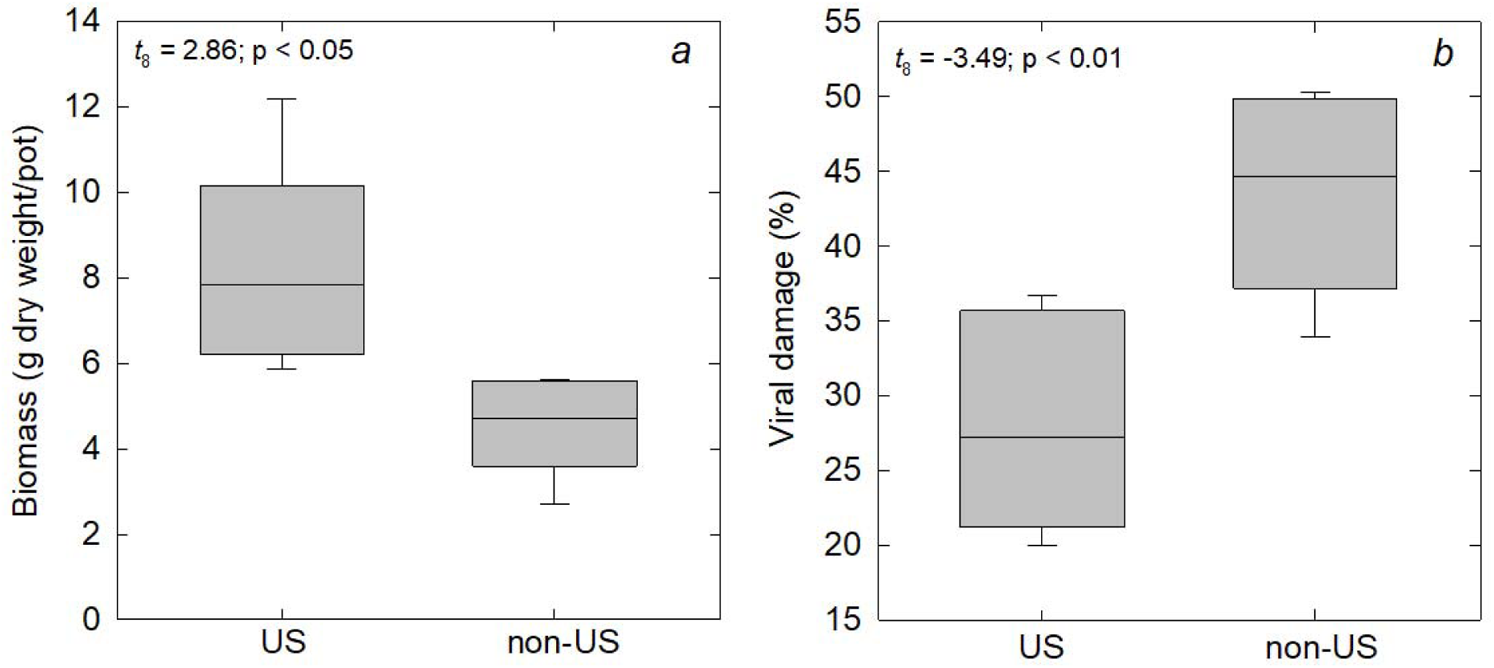
Boxplots of biomass production (a) and leaf viral damage (b) from cultivars domesticated in the United States (US) versus outside the US. Differences between means were assessed using a t-test (t-statistic with 8 degrees of freedom).

## Discussion

Hops have no doubt become a dominant cash crop, feeding into the growing beer industry across the world. With increased global temperatures and corresponding biotic and abiotic effects of climate change affecting growth patterns and yields, we sought to address three important questions about hop biology: (a) are hops globally structured, and if so, can we establish their evolutionary demographic history with genome-wide variants, (b) are rhizospheric microbial communities different between US-based and non-US-based hop cultivars, and if so, are they correlated with soil properties?, and (c) are agronomically relevant traits such as viral damage and biomass production correlated with their domestication history? We address these questions with a combination of population genomic analyses of genome-wide variants, common-garden experiments, morphological, and soil measurements. Our findings indicate (1) significant population structure amongst cultivars by ancestry, with high genetic differentiation amongst cultivars of American origin, (2) significantly low effective population sizes and genetic diversity due to bottlenecks post divergence in American and Noble ancestral cultivars, (3) earliest estimated divergences during domestication indicating a split between common ancestors of contemporary American cultivars and Central European cultivars around 2800 ybp, with the most recent divergence occurring around 600 ybp to yield contemporary English cultivars, (4) significant continued gene flow and crossbreeding between Central European cultivars, Noble, and English strains, (5) significant differences in the rhizospheric microbial communities of American vs non-US-based cultivars, (6) significant differences in agronomically relevant traits such as leaf viral damage and biomass between American and non-US-based cultivars, (7) significant correlation between dissolvable nitrogen in rhizospheric soils and growth rates.

### Global Population Structuring of Hops

Global population structuring of analyzed hop cultivars largely coincides with their known origination and breeding histories. For instance, Hersbrucker Alpha, determined to be the least diverse cultivar, is the result of a clonal selection of the popular landrace Hersbrucker (USDA Hops Cultivar Database), selectively bred for elevated alpha acid levels. Alpha acids are one of two bittering acids that are of importance to brewers and dictate the overall bitterness of the final beer product (Cerenak et al. 2006). Typically, the lower the alpha acid (3-6%), the less bitter the beer, the higher the alpha acid content, the more bitter or ‘hoppy’ the beer. Hersbrucker is a Noble hop which typically imparts pleasant, ‘European style’, aromas and has an alpha acid content of roughly 5-6%. While it was reported by the USDA that Hersbrucker Alpha was not found to display high alpha acid content, the process of clonal selection and intense selective breeding tactics are likely causal to its low estimated genomic diversity. Banner on the other hand is the result of open pollination with Brewer’s Gold, a cultivar that is also the result of open pollination. Open pollination is a breeding tactic used to increase genetic diversity amongst cultivated populations (Rauf et al. 2010). Brewer’s Gold is the ancestor of most of the major high-alpha hops around the world, and as a result has been used extensively in selective breeding. Our results suggest that Banner has overall higher genomic diversity than its progenitor, and therefore has potential for selective open pollination and future breeding purposes with the intention of retaining that high alpha hop trait.

Results of our relatedness analysis further support our population groupings as suggested by ADMIXTURE. Cultivars of Group A from our relatedness analysis shared a high degree of relatedness due to recent outcrossing with wild American hops. For example, Brewer’s Gold and Bullion are both F1 hybrids, resulting from open pollination of a Wild Manitoba (potentially *Humulus lupulus var. lupuloides*) hop. Additionally, several cultivars within Group 1 are known offspring of Brewer’s Gold (Centennial, Aquila, Olympic and Galena) or F2 hybrids (Chelan). All members of Group 2 and 3 are found within their respective ancestry groups (Noble and American respectively), suggesting high degree of relatedness and population structure (due to open pollination within geographical areas, or selective breeding) within each ancestry group.

Both Noble and Central European cultivars are predominantly derived from the Central European region. The Noble ancestry group consists mainly of hops that have been traditionally used in the brewing industry in Europe. Central European hops on the other hand comprise many of the known landrace hops and their descendants. Meanwhile, cultivars of American ancestry share a recent outcrossing event with wild American hops, resulting in F1 and F2 hybrids within this population. Interestingly, American ancestry and English ancestry hops shared similar F_st_ levels as those between Central European/Noble hops. Historically, hops were brought by the English to Massachusetts sometime during the seventeenth century with the first commercial production beginning in 1648 (Korpelainen and Pietiläinen 2021). While our findings suggest that English and American ancestry hops share many genetic similarities, phenotypically these hops have been known to serve different functions in the brewing industry. Many of the cultivars within the English ancestry group are known to impart light, clean, herbal and citrusy notes in the beer they are used to brew, whereas cultivars of the American ancestry group are known to impart more intense notes of earthiness, spice and resin, suggesting that the subtle genetic variation between the two groups may be a result of phenotypic selection for differing aroma profiles.

Many landrace varieties in Group 1 showed 100% assignment to Central Europe. These include Hersbrucker and many of its clonal selections (Hersbucker G, Hersbrucker Alpha, Hersbrucker Redstem, and Hersbrucker 6) as well as Wuerttemberger and Tardif d’ Bourgogne. Additionally, Fuggle N, Yugolding, Styrian and Savinja Golding were also reported as displaying 100% assignment to Group 1 and are all clonal selections of one of the oldest known hops landraces, Fuggle. Fuggle is of English origin, has often been grouped with the ‘Goldings’ such as Styrian, Savinja and Eastwell/Eastkent Golding, due to similar properties such as ‘pleasant’ aroma and similar alpha acid concentrations (5.1%-6%). Additionally, all cultivars assigned to Group 1 are known as ‘aroma’ hops, and in some cases, as dual-purpose hops, serving both as aroma agents as well as bittering hops.

Only four cultivars in Group 2 were estimated to be of “purely” Noble ancestry (Osvald 72yvf, Tettnanger, Nadwislanka and Saazer 38). Three of these four cultivars are known ‘Noble’ hops, renowned for their indicative ‘pleasant’ or ‘continental-style’ aromas. Noble hops have been used in brewing classic European style beers such as lagers or pilsners. Clonally selected progeny of the fourth known Noble hop, Spalter, called “Spalter Select” used in our analyses, clustered with other Noble hops in Group 2.

In Group 3, most cultivars were estimated to have a high degree of admixture between cultivars in the Central European and Noble ancestry groups. Many of the cultivars in Group 3 are of English ancestry and are known ‘bittering’ or dual purpose hops, meaning their primary purpose for cultivation is to serve as a necessary bitterer to counteract the sweetness of the grains used in beer production (Ayabe et al. 2018.

Lastly, Talisman was the only cultivar to present 100% ancestry to the American ancestry group. Talisman is the offspring of the oldest hops cultivar known in North America, Late Cluster, undergoing an open pollination. We suspect that this open pollination may have occurred with some Wild American hops due to sharing similar proportions of ancestry with cultivars such as Brewer’s Gold, which are the result of open pollination with wild American hops (USDA ARS Hops Cultivar database).

In addition to revealing similar topologies to the population structure analyses, our phylogenetic tree was able to untangle some unknown pedigree history for some cultivars. For instance, according to the USDA-ARS Hops Cultivar database, the origin of the German cultivar Hersbrucker and most of the varieties (‘hersbruckeg’, ‘hersbruckeralpha’, ‘hersbrucker6’) were of unknown pedigree, suspected to be relatives of the Czech landrace ‘Saazer’ (USDA-ARS Hops Cultivar database). Our phylogenetic analysis revealed that many of the Hersbrucker varieties (‘hersbrucker’, ‘hersbruckeralpha’ and ‘hersbrucker6’) including the original selection of Hersbrucker (‘hersbruckerg’) was shown to be sister taxa to the French landrace Tardif d’ Bourgogne, supported with an 80% bootstrap value. Additionally, Tardif d’ Bourgogne occurs in the Alsace region of France which lies close to the French/German border, suggesting an eastward migration. Interestingly, this is concordant with a period of emigration of French people into Germany during historical events such as the St. Bartholomew Day massacre (1572), and the French emigration (1789-1815). Our findings suggest a possible connection between human and plant migration.

### The History of Hops Domestication

Using the thousands of simulated demographic models generated by CoalMiner and fastsimcoal28, our top model, and therefore most likely demographic model revealed three key evolutionary time points in the domestication history of hops; 1) divergence between Central European and English cultivars around 623.5 ybp (95% C.I. 629.64 - 648.18 ybp) 2) divergence between Central European and Noble cultivars with Noble hops undergoing a bottleneck event around 2337 ybp (95% C.I. 2352.95 - 2429.61 ybp) and 3) divergence between North American and Central European cultivars about 2799 ybp (95% C.I. 2724.41 - 2799.15 ybp). Interestingly, many of these estimated dates fall in line with the known history of human migration. For instance, during the Neolithic areas many regions around the world began to serve as hotspots for domestication at various time points from as early as 10,000 years ago along the fertile crescent, to more recent times such as that along Eastern North America roughly 3,000 years ago. Our estimated time point of 2799 years ago for the divergence between North American and Central European cultivars falls close to this known timeframe of domestication within North America. Similarly, the estimated time of divergence between Central European and English hops cultivars falls closely within the estimated time window of hops being introduced into the Kent area of England toward the end of the 15^th^ century from Central Europe (History of Hops 2022).

### Molecular ecology of the rhizosphere in US vs non-US hop cultivars

Our study establishes that the rhizospheric microbial community systematically and functionally differs amongst new and world world hop cultivars in a controlled setting, paving the way for additional field-based research into the role of soil microbes in agronomically relevant hop traits like biomass, humulone and co-humulone concentrations, drought tolerance, and viral/viroid resistance. Overall, US based cultivars rhizospheres displayed lower levels of alpha diversity, specifically with lacking nitrogen fixing bacteria, while producing more bacterial gene products (Fig. 5). This observation suggests that fewer, less diverse bacterial strains have likely co-evolved with domesticated new world cultivars, while being more functionally productive. Particularly, US cultivar soils uniquely exhibited the presence of Micrococcales, a genus associated with salt, drought, and alkali resistance across arid soil ecosystems (Sun et al., 2020), Rhizobiales that are associated with nitrogen fixing (Freiberg et al., 1997), and families of methylotrophs. Control soils (potting soil) uniquely exhibited the presence of Ruminococcaceae, a genus of bacteria that are commonly found among the gut microbiota of ruminants and manure (Wang et al., 2013). Meanwhile, non-US cultivar soils showed the presence of another family of Rhizobilaes (Bradyrhizobiaceae) that are also involved in nitrogen fixing (Marcondes de Souza et al., 2014), Myxococcales that produce several important secondary metabolites (Saggu et al., 2023), and interestingly, the presence of predatory bacteria in Vampirococcus genus. While some recent studies have attempted to understand the roles of soil microbes in hop creep (spoilage of hopped beer due to breakdown of grain starch compounds by amylases on the surface of the hop bines - Young et al., 2023), or the role of microbial communities in autotoxicity in hops (Zhang et al., 2011), there is as yet an incomplete understanding of the role of rhizospheric microbial communities in hop agronomic trait variation. For instance, our modeling of plant growth rate and biomass indicates (a) non-dependence of growth and biomass on the presence/absence of nitrogen fixers, and (b) significant dependence of growth and biomass on soil dissolvable nitrogen. However, hops produce several volatile compounds, variable concentrations of which are likely to be affected by above and below ground microbial activity. We contend that more studies will therefore be required to characterize and analyze enzyme functions of microbial enzymes in tandem with bitter acid concentration across hop cultivars. Our study provides a baseline database of operational taxonomic units (OTUs) as observed across domesticated cultivars, which can be expanded in future comparative studies.

**Fig. 5:**
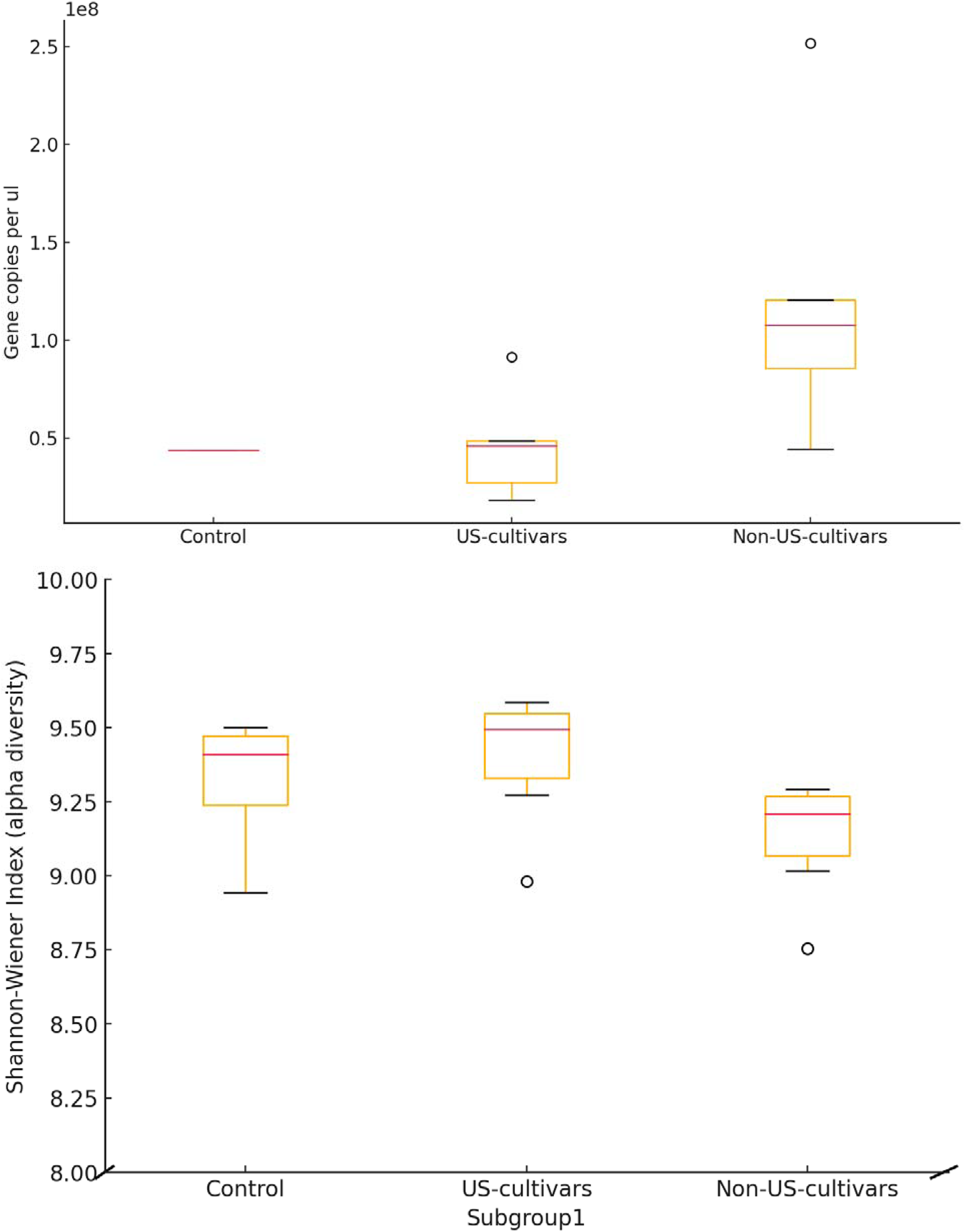
(Top) Boxplots showing absolute abundance, here measured as the total number of 16s rRNA V3-V4 gene copies per ul, and (Bottom) Distribution of Shannon-Wiener (alpha) diversity indices showing the weighted proportional abundances of bacterial species identified in rhizospheric soil samples. “Control” pots were maintained with no hop rhizomes, “Non-US-cultivars” indicate - non-US domesticated hop cultivar pots, and “US-cultivars” showing US-domesticated cultivar pots.

### Agronomic trait variation in US vs non-US hop cultivars

We found that the new world hop varieties had higher rates of biomass production than the European varieties, which is presumably due to the warm growth temperatures associated within the glasshouse during the southern California summer. Hops production is typically restricted to areas that are above 35° north and south latitude (Turner et al. 2011), where climatic conditions such as low temperature during dormancy, mild temperature, and adequate moisture during the growth phase, occur more frequently than in other regions (Rossini et al. 2021). Many of the European varieties originated in northern latitudes, where growing season temperatures are likely lower than those experienced in the southwest US. For example, Mozny et al. (2009) found that increasing temperatures, above 21°C in the growing season, resulted in decline in Saaz hop yield. A follow up study using simulated data by Mozny et al. (2023) found that going into the latter half of the 21st century, cultivars grown in dominant hops growing regions throughout Germany will continue to see a decline in hops production as a result of increasing temperatures. In growth trials in the Mediterranean Basin, which has a similar climate to southern California, Cascade (a U.S. variety) had higher rates of growth and cone production than the other European varieties (Ruggeri et al. 2018). Similarly, rates of C assimilation for US hop strains have a temperature optimum of 21-39°C (Eriksen et al. 2020), which are higher than the optimum temperatures for European varieties such as Saaz (Mozny et al., 2009). Thus, while the temperature sensitivity of hops growth is not known for the strains tested here, we speculate that the European varieties, by virtue of the origin of their domestication, may have had a lower growth temperature optimum than the US varieties.

We also found that the U.S. hop varieties had lower viral damage than the European varieties, which is likely due to (a) selective breeding for viral resistance among new world cultivars, and potentially (b) lack of genetic diversity for viral resistance among the European cultivars utilized in the greenhouse experiment. Many of the European varieties are closely related and have very little genetic variation compared to the U.S cultivars (Korpelainen and Pietiläinen 2021), which increases their susceptibility to certain hops viruses and viroids (Pethybridge et al. 2008). Many viruses that are common in hops, such as the hop mosaic virus, hop latent virus, Apple Mosaic virus, may have had a European origin (Pethybridge et al. 2008), which may contribute to the higher prevalence in the European strains studied here. Furthermore, if growth conditions were warmer than their temperature optimum, it is likely that temperature and/or drought stress would enhance viral damage to strains that are more susceptible to viral infection (Mozney et al. 2009).

## Conclusion

In the short period of time since the onset of their domestication, hops have been utilized extensively in several applications, leaving the greatest mark on the brewing industry. Selective breeding tactics across the globe have led to subtle genetic differences between the nearly 250 developed cultivars, leaving patterns of population structuring and specific microbial associations. Genome-wide SNP data allowed for the detection of substructuring within Central European cultivars, revealing patterns in agronomic trait selection. Future studies should therefore focus on identifying genomic loci associated with strong global patterns of differentiation and linking them to agronomic traits to benefit future crop improvement efforts. Additionally, follow up studies aiming to understand the above and below ground dynamics between hops and their soil microbial communities show potential for revealing key microbial associations that may be responsible for some cultivars’ abilities to flourish outside of their native ranges. Going into the latter half of this century, this knowledge will be ever so imperative, especially with many of our cash crops (hops included) facing serious challenges under global climate change.

## Data Availability

All data summaries and pipelines utilized in data analyses can be found on the public Github repo: https://github.com/raywray/hops_pipeline.git. SNP data analyzed in this manuscript are property of the USDA-ARS and can be requested from the USDA-ARS.

## Acknowledgments

This work was supported by USDA-HSI:2022-77040-38529 to PI Sethuraman and co-PIs Vourlitis and Jancovich, USDA-REEU: 2017-06423 to PI Vourlitis and co-PI Sethuraman, and NSF CAREER: 2147812 to PI Sethuraman. AMA was supported via Genetics Society of America, California Agricultural Institute NextGen grants. TM was supported via a CSUBIOTECH grant. All computations were performed on the *mesxuuyan* cluster at San Diego State University, which was supported by startup funds and NSF ABI: 1564659 to AS. We thank members of the Sethuraman Lab for helpful suggestions on early versions of this manuscript. All SNP data analyzed in this manuscript are subject to the Data Transfer Agreement between the USDA-ARS and San Diego State University, dated January 5, 2022. We graciously acknowledge Dr. John Henning (USDA-ARS) for generating and sharing these data.

## Supplementary Tables and Figures

**Fig. S1:**
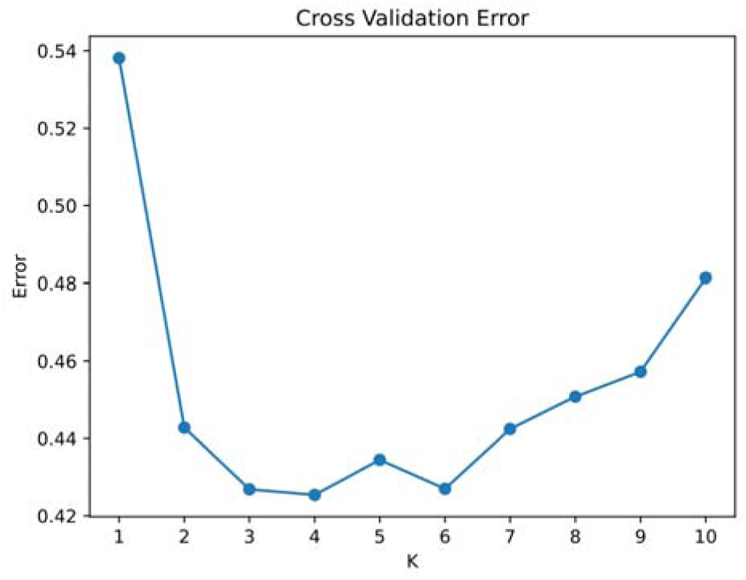
Cross validation (CV) error estimates from replicate runs of ADMIXTURE (Alexander et al., 2009) across bi-allelic SNPs, indicating that the least CV error and therefore most “optimal” clustering of all hop cultivars analyzed was achieved at K = 4 subpopulations. Based on population assignment, these 4 subpopulations were thereon labeled as “Noble”, “English”, “American”, and “Central European” based on their known historical provenances.

**Fig. S3:**
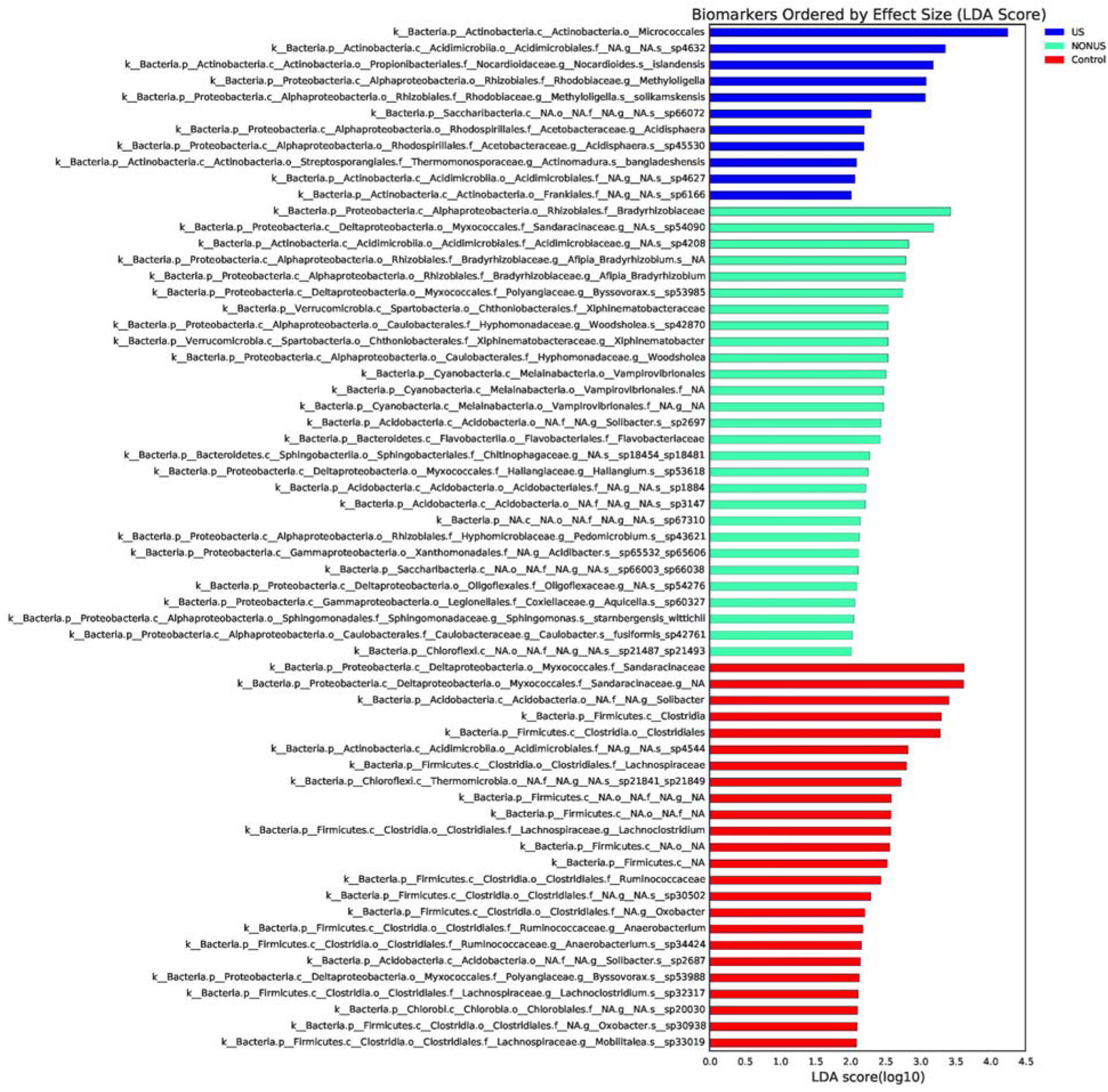
LDA plot generated using LEfSe analyses to identify bacterial taxonomic distributions that are significantly different among “Control”, “US” and “Non-US” based cultivar rhizospheric soils, here weighted by effect size and showing only bacterial taxa that have an LDA score > 2.0.

**Fig. S4:**
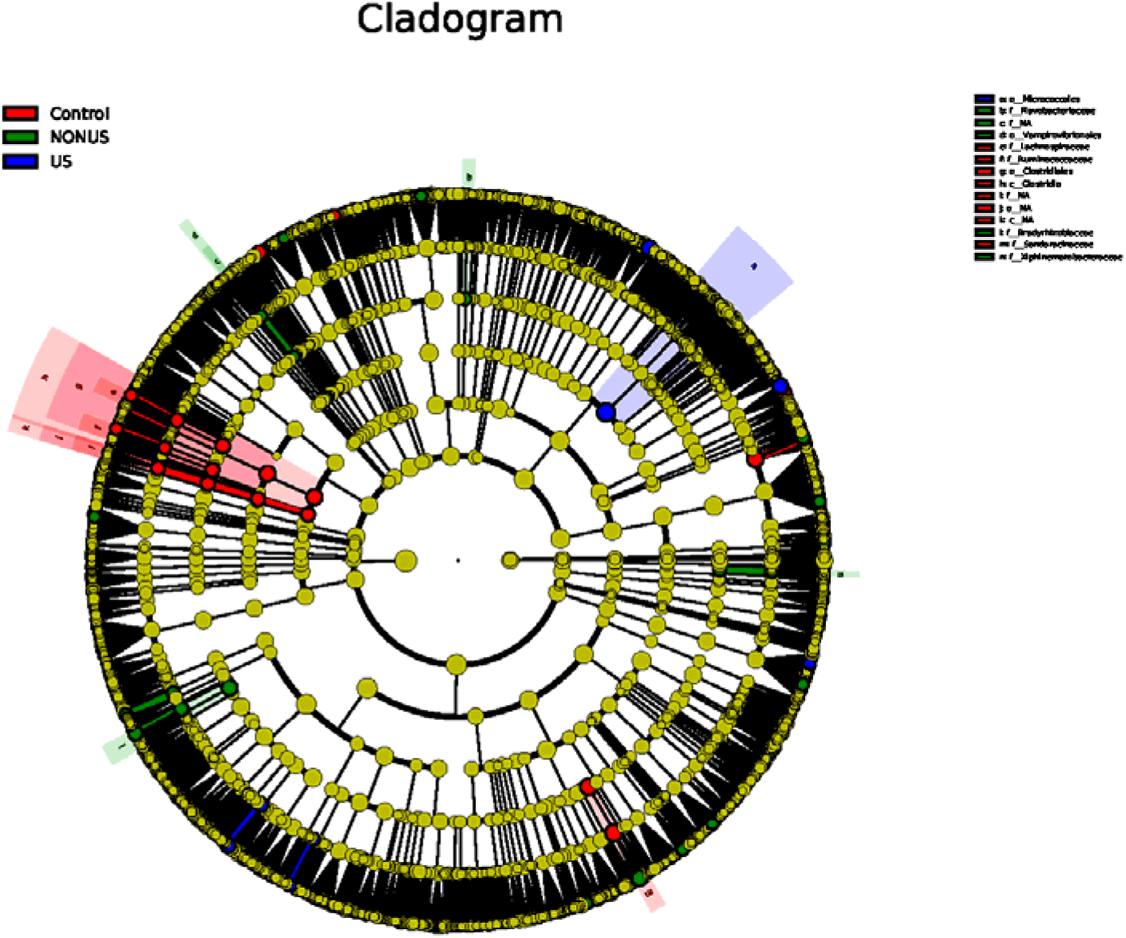
A cladogram of identified bacterial taxa, generated using LEfSe analyses to show bacterial taxonomic distributions that are significantly different among “Control”, “US” and “Non-US” based cultivar rhizospheric soils, here indicating only taxa that are uniquely present in each grouping with a p value < 0.05.

**Table S1:** Variance Inflation Factor analysis of model predictors mean dissolved nitrogen ”mean_DIN”, genus relative abundance “genus_rel_abun” and nitrogen fixing capability “n_fixing”. All model predictors show adjusted generalized variance inflation factor <10 indicating no multicollinearity exists among predictors.

| predictor | GVIF | Df | Adjusted GVIF |
| --- | --- | --- | --- |
| mean_DIN | 1.000204 | 1 | 1.000102 |
| genus_rel_abun | 1.040676 | 1 | 1.020135 |
| n_fixing | 1.040472 | 3 | 1.006634 |

**Table S2:** Assessment results of Quasi-Poisson GLM. Average dissolved nitrogen, “mean_DIN” shows a significant positive effect on average relative growth rate.

|  | Estimate | Std Error | T value | P value |
| --- | --- | --- | --- | --- |
| (intercept) | 0.929370 | 0.024155 | 38.475 | <2e-16 * |
| mean_DIN | 0.235590 | 0.019089 | 12.342 | <2e-16 * |
| genus_rel_abun | 0.002423 | 0.021359 | 0.113 | 0.910 |
| n_fixingYes | 0.001006 | 0.048996 | 0.021 | 0.984 |

**Fig. S5:**
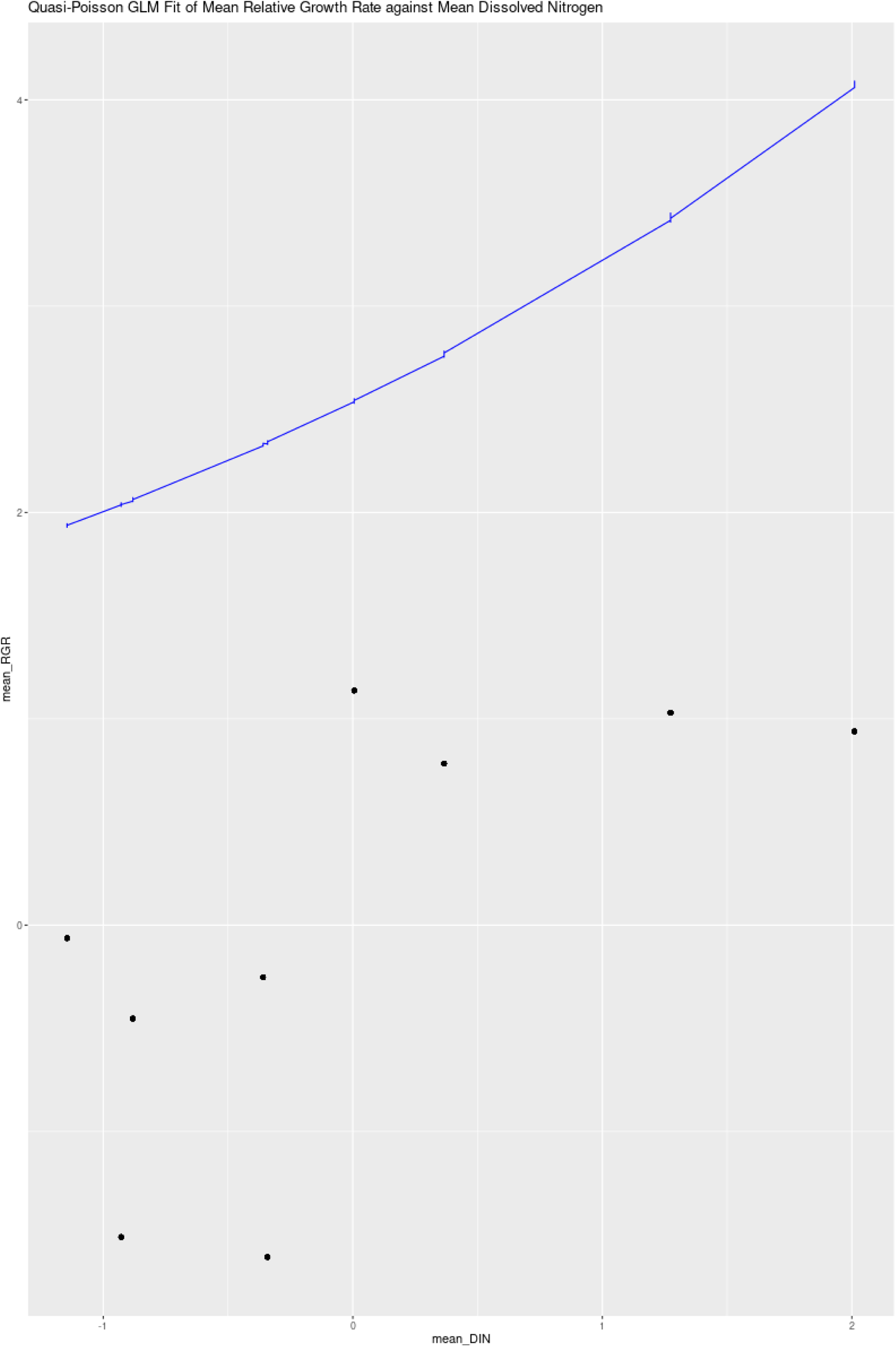
A Predicted vs observed plot of average relative growth rate against average dissolved nitrogen content.

**Fig. S6:**
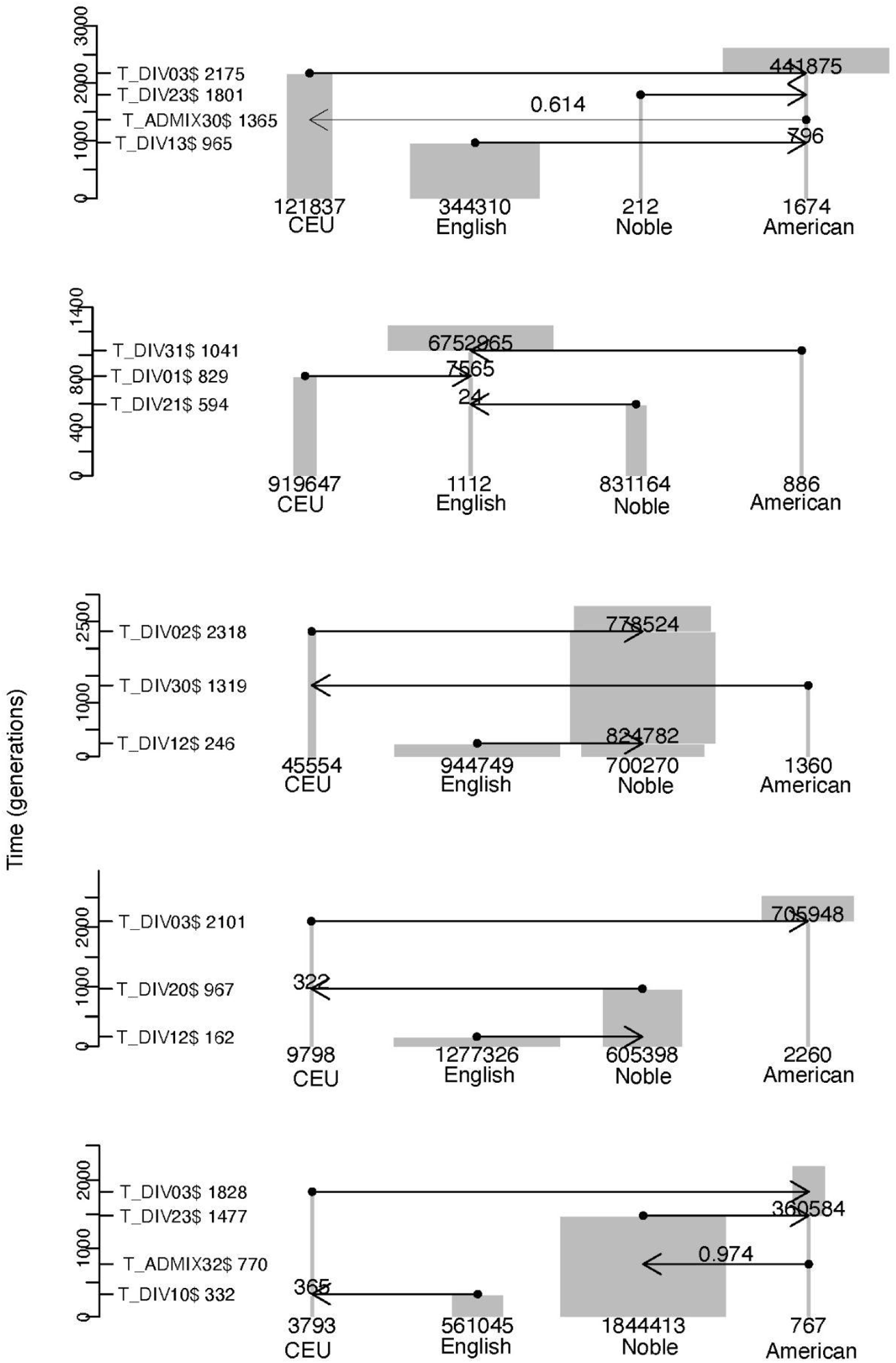
Next top 5 (after best model, shown in Fig. 2) predicted evolutionary demographic models using fsc28 + CoalMiner. Group 1 = Central European Ancestry, Group 2 = Noble Ancestry, Group 3 = English Ancestry, Group 4 = American Ancestry.

**Table S3:** Log likelihoods estimated from top 10 runs of fsc28.

| Model | Log Likelihood |
| --- | --- |
| model899 | -17619.616 |
| model211 | -17713.841 |
| model754 | -17729.413 |
| model635 | -17888.942 |
| model449 | -17900.052 |
| model550 | -17921.002 |
| model640 | -17923.157 |
| model541 | -17936.673 |
| model675 | -17974.329 |
| model778 | -17992.77 |

**Fig. S7:**
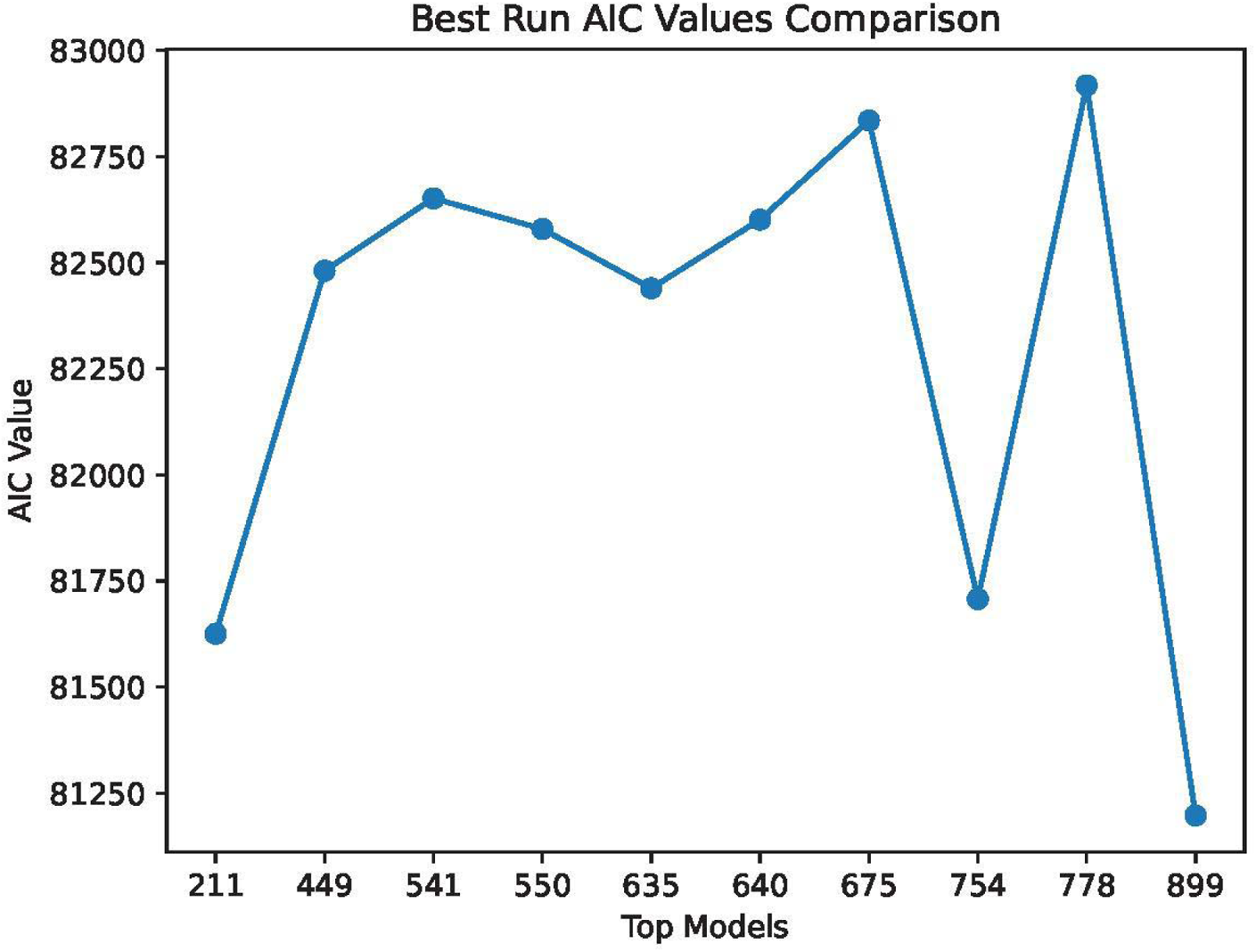
AIC values determined from top ten models using fsc28.

**Table S4:**
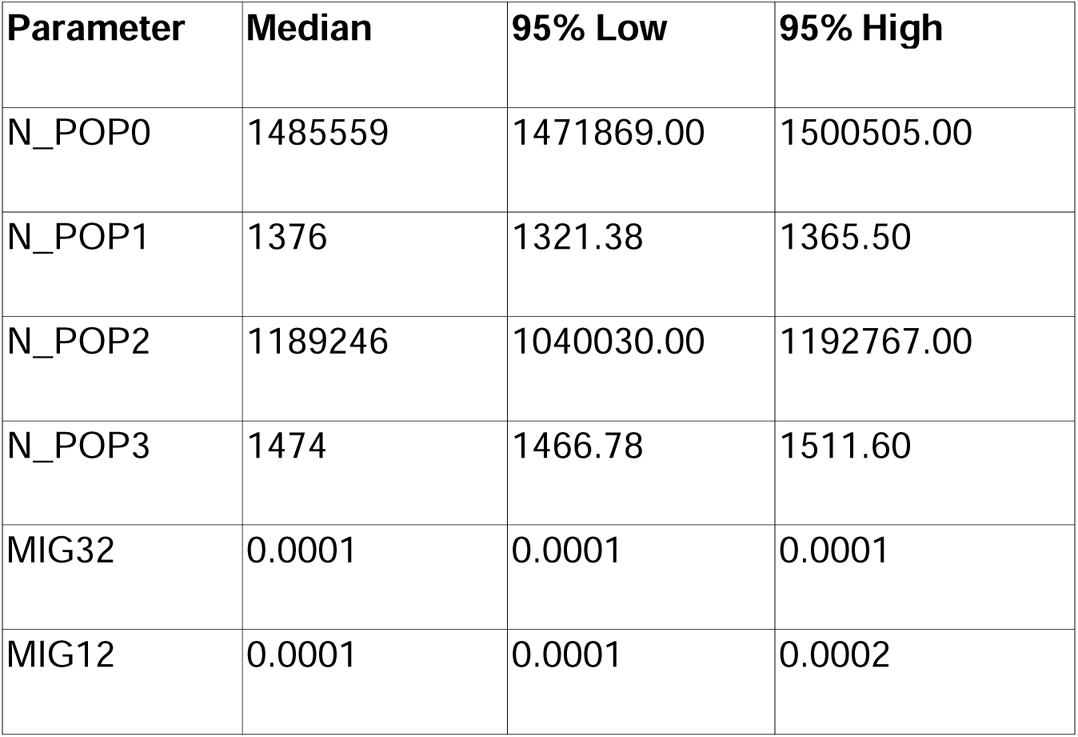
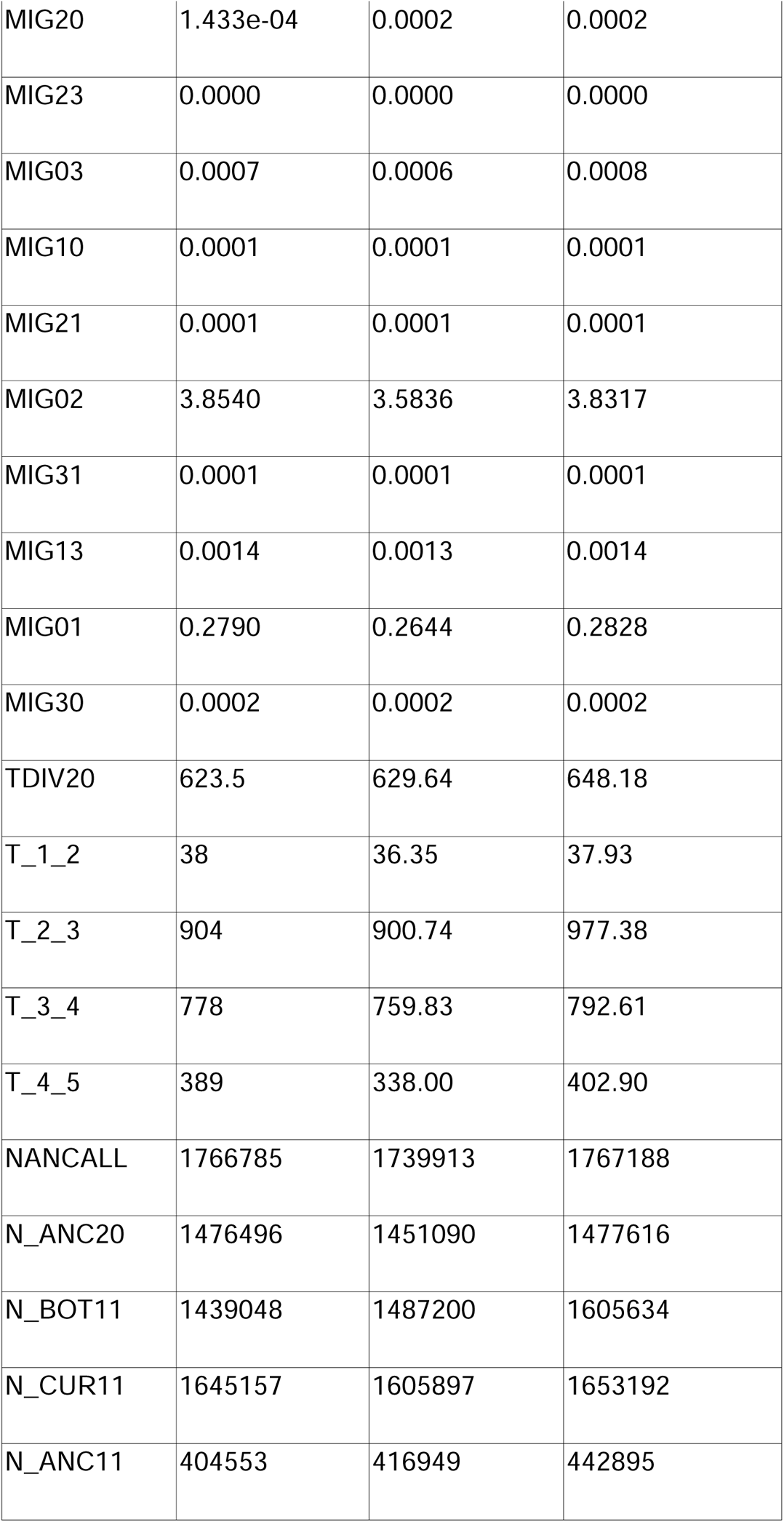
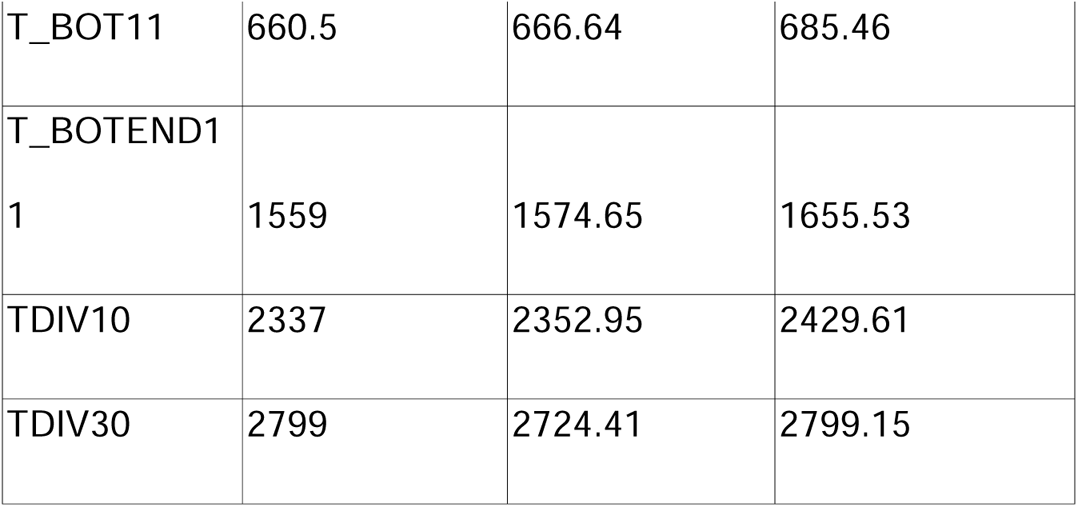
Median, and 95% confidence intervals around demographic parameter estimates for the top fsc28 model.

